# TgCentrin2 is required for invasion and replication in the human parasite *Toxoplasma gondii*

**DOI:** 10.1101/491316

**Authors:** Jacqueline M. Leung, Jun Liu, Laura A. Wetzel, Ke Hu

**Affiliations:** Department of Biology, Indiana University, Bloomington, IN, 47405, USA; Current address: Department of Molecular and Cellular Biology, University of California, Berkeley, CA, 94720, USA

## Abstract

Centrins are EF-hand containing proteins ubiquitously found in eukaryotes and are key components of centrioles/basal bodies as well as certain contractile fibers. We previously identified three centrins in the human parasite *Toxoplasma gondii*, all of which localized to the centrioles. However, one of them, TgCentrin2 (CEN2), is also targeted to structures at the apical and basal ends of the parasite, as well as to annuli at the base of the apical cap of the membrane cortex. The role(s) that TgCentrin2 plays in these locations was unknown. Here we report the functional characterization of TgCentrin2 using a conditional knockdown method that combines transcriptional and protein stability control. The knockdown resulted in an ordered loss of TgCentrin2 from its four compartments, due to differences in incorporation kinetics and structural inheritance over successive generations. This was correlated with a major invasion deficiency at early stages of *TgCentrin2* knockdown, and replication defects at later stages. These results indicate that TgCentrin2 is incorporated into multiple cytoskeletal structures to serve distinct functions in *T. gondii* required for parasite survival.

## INTRODUCTION

Centrin-like proteins were first discovered in the ciliated protozoan *Zoothamnium geniculatum* as a component of the spasmoneme, an organelle that contracts in response to calcium binding (Amos et al., 1975). Subsequently, centrins were found to be components of the centrioles/basal bodies and to play an important role in regulating centrosomal duplication, and flagellar biogenesis and function in the cells of many animals and protozoa (Salisbury, 1995; Salisbury, 2007; Sanders and Salisbury, 1994; Wright et al., 1985).

We previously identified three centrins in *Toxoplasma gondii*, a human parasite that causes devastating toxoplasmic encephalitis in immunocompromised individuals and unprotected fetuses. As with canonical centrins in other systems, TgCentrins 1 and 3 (CEN1 and 3) are predominantly localized to the centrioles. Although TgCentrin2 (CEN2) shares a high degree of sequence similarity with CEN1 and 3 (Figure 1A), its localization is remarkably different (Hu et al., 2006). In addition to the centrioles, ectopically expressed CEN2 tagged with eGFP also localized to three other structures: a ring-shaped complex at the apex of the parasite (the preconoidal rings), a capping structure at the basal end (the basal complex), as well as ~5-6 peripheral annuli located approximately one-quarter of a parasite length below the apex (Hu et al., 2006) (Figure 1B). The location of the CEN2 annuli coincides with the boundary between the apical cap and the rest of the parasite membrane cortex, as marked by the ISP proteins (Beck et al., 2010).

**Figure 1.**
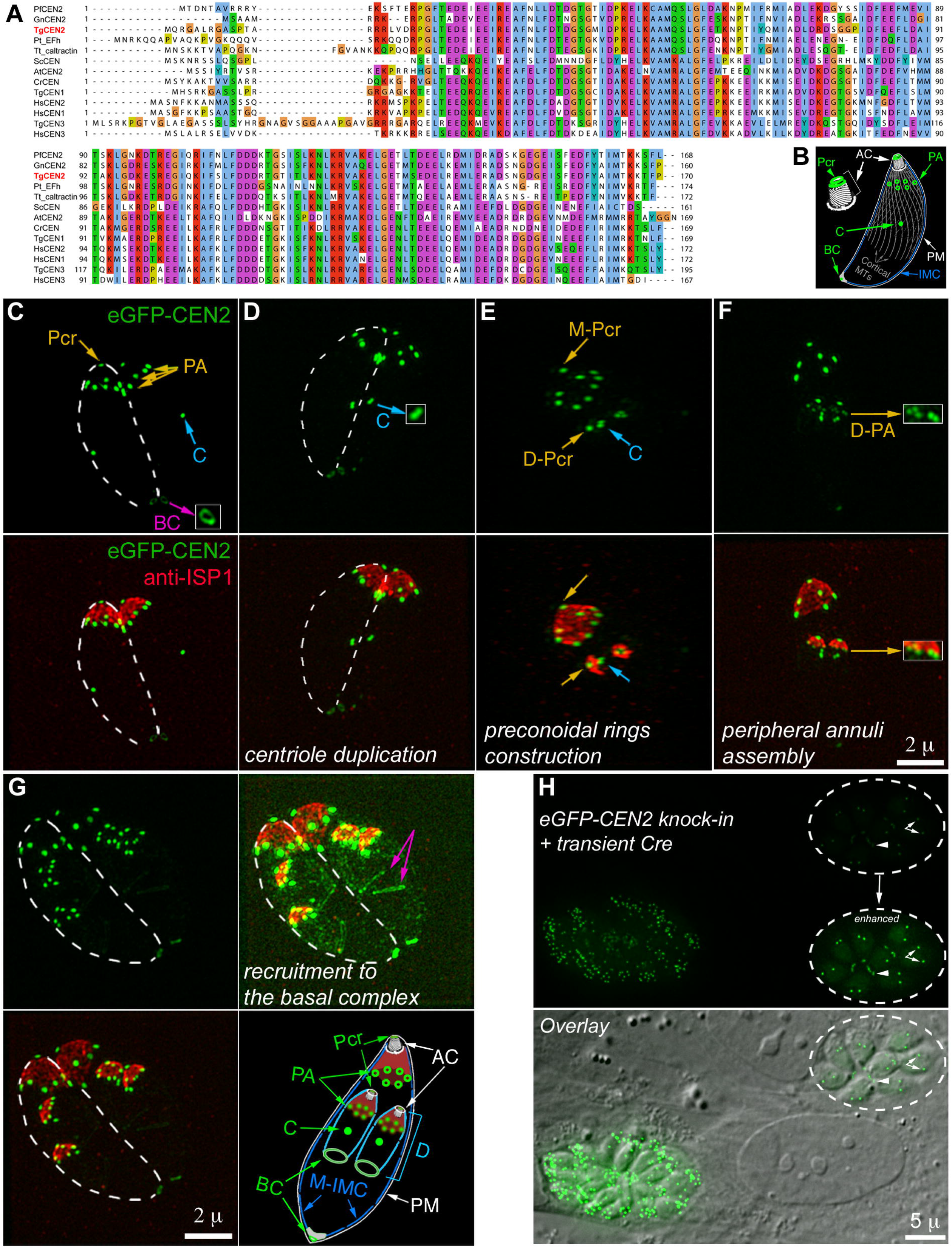
Phylogenetic analysis of selected centrin homologs and the localization of TgCentrin2 (CEN2) in *Toxoplasma gondii* (A) Alignment of TgCentrin1-3 (TgCEN1-3) and selected centrin homologs in other organisms using the multiple alignment program MUSCLE. PfCEN2 XP_001348617.1 (GenBank), GnCEN2 XP_011129982.1, TgCEN2 TGGT1_250340 (ToxoDB), Pt_Efh XP_001441649.1, Tt_caltractin XP_001023350.1b, ScCEN NP_014900.3, AtCEN2 NP_190605.1, CrCEN XP_001699499.1, TgCEN1 TGGT1_247230, HsCEN2 NP_004335.1, HsCEN1 NP_004057.1, TgCEN3 TGGT1_260670, HsCEN3 NP_004356.2, Pf: *Plasmodium falciparum*; Gn: *Gregarina niphandrodes*; Tg: *Toxoplasma gondii*; Pt: *Paramecium tetraurelia*; Tt: *Tetrahymena thermophila*; Sc: *Saccharomyces cerevisiae*; At: *Arabidopsis thaliana*; Cr: *Chlamydomonas reinhardtii*; Hs: *Homo sapiens*. (B) Cartoon of an interphase parasite, in which CEN2-containing structures are highlighted in green. Pcr: preconoidal rings; PA: peripheral annuli; C: centrioles; BC: basal complex; AC: apical complex; PM: plasma membrane; IMC: inner membrane complex; Cortical MTs: cortical microtubules. (C-G) Projections of 3D-SIM images of *eGFP-CEN2* knock-in parasites at different stages of the cell cycle labeled with a mouse anti-ISP1 antibody. Insets (2X) in C, D, and F are contrast enhanced and include regions indicated by the arrows. Dashed lines in C, D, and G indicate the approximate outline of one of the two parasites in the same vacuole. The cartoon in G highlights the localization of CEN2 (green) and ISP1 (red) with respect to the inner membrane complex (IMC) and the plasma membrane. “M-”: mother structures. “D-”: daughter structures. D: daughter. Other abbreviations are the same as in (B). Green: eGFP-CEN2; Red: anti-ISP1. Scale bar = 2 μm. (H) Fluorescence (top) and fluorescence/DIC overlay (bottom) images of *eGFP-CEN2* knock-in parasites ~48 h after transfection with a plasmid expressing Cre recombinase. There are two vacuoles in the field. Dashed circles indicate the vacuole in which eGFP-CEN2 expression has decreased significantly. Inset (1X) is contrast enhanced to visualize residual eGFP-CEN2 signal in the centrioles (arrows) and basal complex (arrowheads) of the parasites in this vacuole. Scale bar = 5 μm.

We were interested in understanding what function(s) CEN2 plays in the four distinct structures that it targets. After many failed attempts with established gene manipulation approaches to delete the *CEN2* gene or downregulate its expression, we knocked down *CEN2* expression in *Toxoplasma* using a dual-regulation approach that combines anhydrotetracycline (ATc)-mediated transcription suppression and ddFKBP-mediated protein degradation. We discovered that CEN2 was depleted from its four locations with different kinetics. The loss of CEN2 from the two more apically located structures, the preconoidal rings and peripheral annuli, occurred earlier, followed by significant CEN2 depletion from the centrioles and basal complex. This was correlated with a major inhibition of parasite invasion evident at early stages of *CEN2* knockdown followed by replication defects that developed at later stages. This suggests that CEN2 is critical for multiple aspects of the parasite lytic cycle and that its associated structures comprise potential targets for anti-parasitic measures.

## RESULTS

### The localization of CEN2 in eGFP-CEN2 knock-in parasites

In the previous studies, the localization of CEN2 was characterized by ectopic expression. To confirm its localization pattern, we replaced the endogenous *CEN2* gene with *eGFP-CEN2* using homologous recombination. The resulting *eGFP-CEN2* knock-in parasite line shows that CEN2 is localized to the preconoidal rings, peripheral annuli, centrioles, and basal complex (Figure 1B-G), the same pattern as ectopically expressed eGFP-CEN2 (Hu, 2008; Hu et al., 2006). Figure 1C-G includes images of parasites at different stages of cell division – a “born-within” type of replication (Hu et al., 2002; Nishi et al., 2008; Sheffield and Melton, 1968) – during which the cortical cytoskeleton of the daughter parasite is built inside the mother. Of the CEN2-containing structures, duplication of the centrioles occurs first (Figure 1D, inset), followed by construction of the preconoidal rings (Figure 1E), and then the peripheral annuli, which appear after the ISP1 cap is assembled (Figure 1F&G). Recruitment of CEN2 to the daughter basal complex becomes evident only at a late stage of daughter assembly (Figure 1G).

### Unsuccessful attempts to generate CEN2 knockout and knockdown mutants using several established methods

In the knock-in line, the *eGFP-CEN2* coding sequence and the *HXGPRT* drug selection cassette are flanked with *LoxP* sites, which allows for the excision of this locus upon transfection with Cre recombinase-expressing plasmids. Indeed, significant loss of eGFP-CEN2 fluorescence was observed in many parasites ~48 h after Cre transfection (Figure 1H). However, numerous attempts to isolate *CEN2* knockout clones failed. We also failed to recover *CEN2* knockout clones using a CRISPR-Cas9 based gene deletion strategy (data not shown), suggesting that the complete loss of *CEN2* results in lethality. This result is consistent with the fitness score for CEN2 (−4.4) obtained by (Sidik et al., 2016).

The functions of many presumed essential genes have been characterized using the anhydrotetracycline (ATc)-based knockdown method (Meissner et al., 2001; Meissner et al., 2002; Mital et al., 2005). Therefore, we attempted to regulate *CEN2* expression by generating a line in which the expression of *mAppleFP-CEN2* was driven by the TetO7Sag4 promoter and could be downregulated with ATc, which binds to the tetracycline-inducible transactivator (TATi) to inhibit transcription initiation by the promoter. We found that while the ATc treatment significantly downregulated *mAppleFP-CEN2* expression in the initial passages, the ATc control of *CEN2* expression waned considerably over time (data not shown), indicating that ATc-based transcriptional control alone is not sufficient to maintain robust regulation of *CEN2* expression.

### Combining transcriptional and translational controls allow for more robust and stable knockdown of CEN2

To achieve tighter regulation of CEN2 expression, we combined the ATc transcriptional regulation system with translational control. Specifically, the *eGFP-CEN2* knock-in parasites were stably transfected with a plasmid that contains a *ddFKBP-mAppleFP-CEN2* expression cassette controlled by the TetO7Sag4 promoter, as well as an expression cassette for TATi *(eGFP-CEN2* knock-in: *ddFKBP-mAppleFP-CEN2* cKD parasites, referred to as “KI:cKD” for simplicity). Upon excision of *eGFP-CEN2* from the endogenous locus by transient Cre recombinase expression, clones (*ddFKBP-mAppleFP-CEN2* cKD parasites, referred to as “cKD”) were isolated and maintained in the presence of Shield-1 (Shld1), which binds to and stabilizes the ddFKBP domain, thus blocking degradation of ddFKBP-mAppleFP-CEN2 (Figure 2). The cellular level of ddFKBP-mAppleFP-CEN2 from the TetO7Sag4 promoter therefore can be controlled not only with ATc (regulating transcription), but also with Shld1, the absence of which leads to proteasome-mediated degradation of the fusion protein (Banaszynski et al., 2006; Herm-Gotz et al., 2007). Figure 3A shows that ddFKBP-mAppleFP-CEN2 correctly localized to all of the CEN2-containing structures, and downregulation of ddFKBP-mAppleFP-CEN2 expression was achieved by withdrawing Shld1 from the culture and adding ATc (-Shld1/+ATc) to a final concentration of 270 nM. To measure the kinetics of CEN2 loss, images of vacuoles that contained two to eight parasites were captured after 12 to 120 h of Shld1 withdrawal and ATc treatment, and mAppleFP fluorescence signals in the preconoidal rings, peripheral annuli, centriole(s), and basal complex were quantified. While Shld1 withdrawal alone (-Shld1/-ATc) resulted in a modest reduction of the mAppleFP-CEN2 signal in the preconoidal rings and annuli, CEN2 localization in these structures became undetectable twelve hours after Shld1 withdrawal and ATc addition (-Shld1/+ATc) (Figure 3B). The depletion kinetics for CEN2 in the centrioles and the basal complex were very different. Withdrawal of Shld1 alone (-Shld1/-ATc) did not result in a significant decrease of the mAppleFP signal in these structures. After 12 h of -Shld1/+ATc treatment, the level of mAppleFP was ~42 ± 2% (centrioles) and 54 ± 5% (basal complex) of the fluorescence in the corresponding structures in cKD parasites without downregulation [Figure 3B, “Baseline” (+Shld1/-ATc)], and ~12 ± 1% (centrioles) and ~10 ± 3% (basal complex) by 48 h. This low level of mAppleFP-CEN2 signal remained detectable even after 120 h of ATc treatment (Figure 3B). As it is unlikely that a protein can persist for ~15-20 generations of the parasite [The parasite doubling time has been estimated to be ~6-8 h (Black and Boothroyd, 2000)], this lingering signal might reflect a residual level of continuing CEN2 synthesis. We observed a similar ordered and differential loss of eGFP-CEN2 from the four compartments in our attempts to generate *CEN2* knockout mutants using the Cre-LoxP based method (Figure 1H).

**Figure 2.**
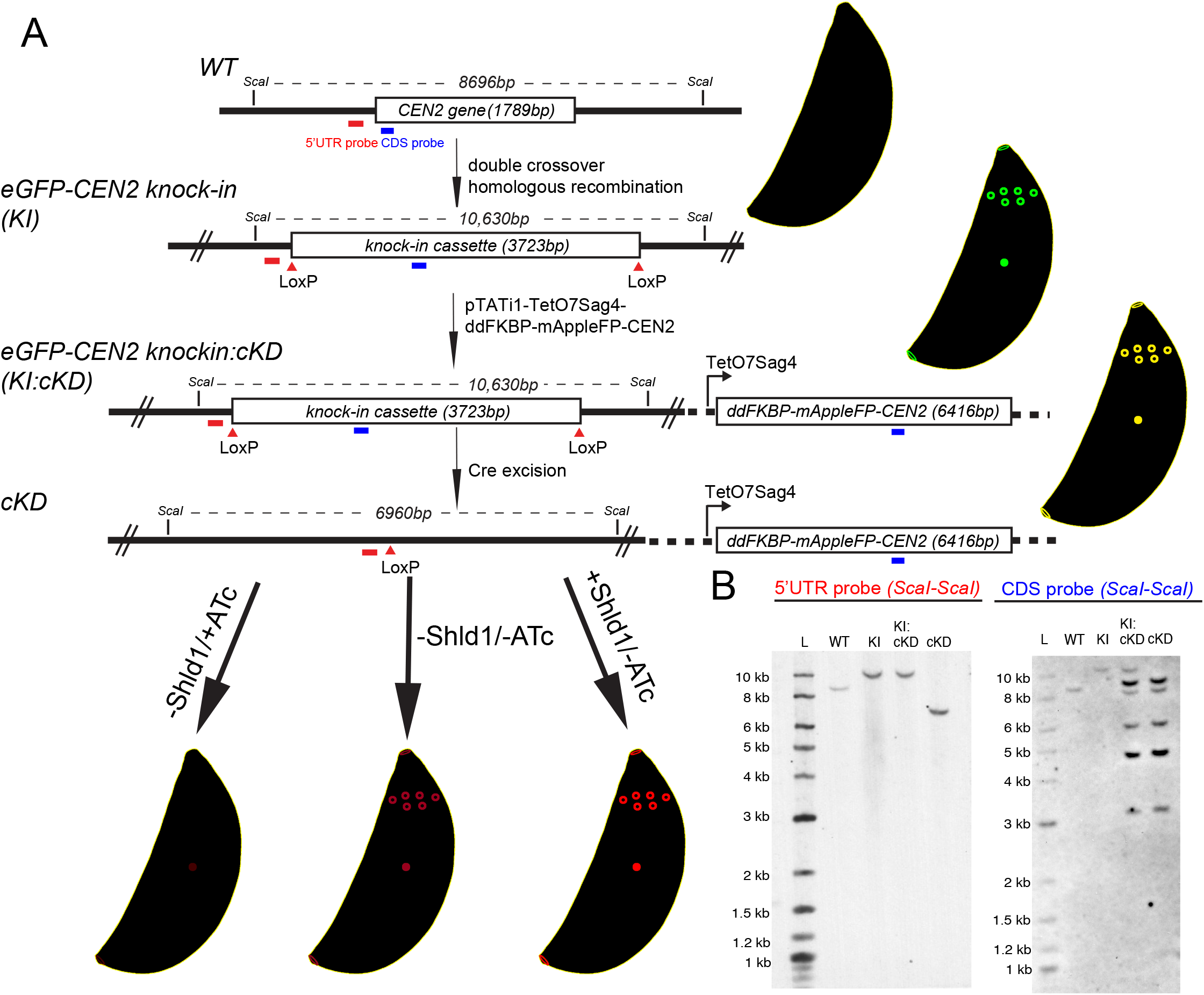
Generation of the *CEN2* knock-in and cKD parasite lines. (A) Schematic for generating *eGFP-CEN2* knock-in (KI), *eGFP-CEN2* knock-in:cKD (KI:cKD) and cKD parasites and Southern blotting strategy. Positions of the *Sca*I restriction sites, CDS probe (blue) and the probe annealing upstream of the *CEN2* coding sequence (5’UTR probe, red) used in the Southern blotting analysis and the corresponding expected DNA fragment sizes are indicated as shown. (B) Southern blotting analysis of the *CEN2* locus in RHΔ*hx* parasites (WT), KI, KI:cKD, and cKD parasites. For the CDS probe, the expected parasite genomic DNA fragment sizes after *Sca*I digestion are 8696 base pairs for the WT (*i.e*., wild-type *CEN2* locus), 10,630 base pairs for the *eGFP-CEN2* cassette in the KI and KI:cKD parasites, and variable for the multiple, random integrations of the cKD plasmid (pTATi1-TetO7Sag4-ddFKBP-mAppleFP-CEN2) since the fragment sizes would depend on where the plasmid integrated relative to *Sca*I sites in the parasite genome. The expected DNA fragment sizes for the upstream probe are the same as for the CDS probe for the WT, KI, and KI:cKD parasites, and 6960 base pairs for the cKD parasites after Cre excision of the eGFP-CEN2 cassette. L: ladder.

**Figure 3.**
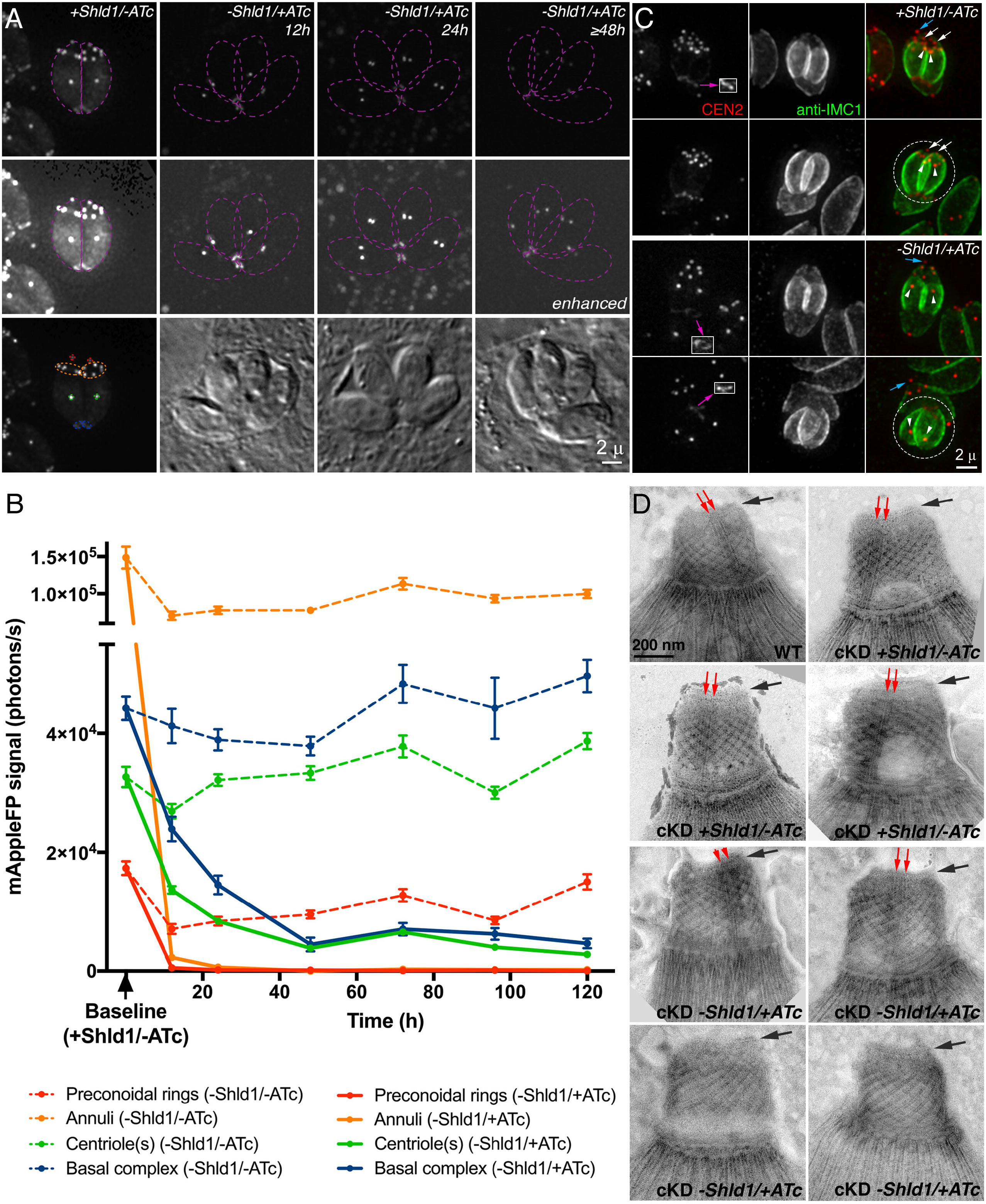
Quantification of CEN2 downregulation in cKD parasites, and ultrastructural analysis of the cytoskeletal apical complex in *CEN2* depleted parasites. (A) Representative fluorescence and DIC images of cKD parasites in two to four parasite vacuoles cultured in the presence of Shld1 (+Shld1/-ATc), or 12, 24, and 48 h after withdrawal of Shld1 and addition of ATc (-Shld1/+ATc). The image of the vacuole cultured in -Shld1/+ATc medium for 48 h is representative of vacuoles imaged at subsequent timepoints (*i.e*., 72, 96, and 120 h -Shld1/+ATc). Individual parasites are outlined in purple, dashed lines. The image of the cKD +Shld1/-ATc parasites is additionally marked with dashed lines to mark the four compartments that CEN2 targets: the preconoidal rings (red), peripheral annuli (orange), centrioles (green), and basal complex (blue). Images in each row were acquired and processed under the same conditions. Images in the middle row were highly contrast enhanced to display low mAppleFP-CEN2 signal in the parasites. Scale bar = 2 μm. (B) Intensity measurements of mAppleFP-CEN2 in the preconoidal rings (red), peripheral annuli (orange), centrioles (green) and basal complex (blue) in cKD parasites 12-120 h after -Shld1/-ATc (dashed lines) or -Shld1/+ATc treatment (solid lines). Baseline: average mAppleFP intensity of individual CEN2 containing structures in cKD parasites cultured in +Shld1/-ATc. Note that the two segments of the y-axis are scaled differently to facilitate better visualization of the lower photon counts in the later timepoints. Error bars: standard error. (C) Fluorescence images of cKD parasites cultured in the presence of Shld1 (+Shld1/-ATc) or treated for 14 h with ATc and no Shld1 (-Shld1/+ATc), in single parasite vacuoles before or undergoing the first round of cytokinesis post-invasion. Dashed circles indicate daughters emerging from the mother parasite. The blue and white arrows indicate the preconoidal rings in the maternal cytoskeleton and newly assembled daughters, respectively, and the white arrowheads indicate daughter centrioles. The daughter basal complexes indicated by the purple arrows are included in the insets (1X, contrast enhanced). Scale bar = 2 μm. (D) Negative staining of whole mount, detergent-extracted parasites to visualize the apical cytoskeletal structure of wild-type parasites (WT), cKD parasites cultured in the presence of Shld1 (cKD, +Shld1/-ATc) or treated for 144 h with ATc and no Shld1 (cKD -Shld1/+ATc). The structure of the preconoidal rings (black arrows) appears to be normal after CEN2 depletion, but the intra-conoid microtubules (red arrows) are not detectable in some CEN knockdown parasites (cKD, -Shld1/+ATc, bottom row). Scale bar = 200 nm.

### The role of structural inheritance in the differential depletion of CEN2 from the four compartments

As shown in Figure 1D-G, the daughter cortical cytoskeleton is built anew inside the mother during the replication of *Toxoplasma* and other apicomplexan parasites, but some cytoskeletal structures, such as the centrioles, are replicated and inherited from the mother (Hu, 2008; Hu et al., 2002). The kinetics of depletion for a protein component from a structure are therefore affected by the synthesis of new protein, the dissociation and degradation rates of pre-existing protein, “dilution” of a finite amount of protein over successive rounds of cellular replication, as well as whether the structure itself is synthesized *de novo* or inherited from the mother. To determine the effect of structural inheritance on CEN2 depletion from the four compartments during knockdown, cKD parasites maintained in Shld1 were allowed to invade, Shld1 was then further maintained, or withdrawn and 270 nM ATc was added to downregulate ddFKBP-mAppleFP-CEN2 for 14 h. Vacuoles containing a single parasite before or undergoing cytokinesis (*i.e*. before completion of the first round of replication post-invasion) were examined. In contrast to cKD parasites maintained in Shld1 where the mAppleFP signal was prominent in all CEN2-containing structures (Figure 3C, +Shld1/-ATc), the ATc-treated parasites assembled daughters with CEN2 that was invariably undetectable in the preconoidal rings and peripheral annuli but prominent in the duplicated centrioles and the basal complex (Figure 3C, -Shld1/+ATc). In some instances, the daughters emerged in an orientation that permitted observation of mAppleFP signal still present in the preconoidal rings and peripheral annuli of the disintegrating mother cytoskeleton (Figure 3C, -Shld1/+ATc, bottom row), strongly suggesting that the maternal CEN2 in these structures was not reused to build the corresponding structures in the daughters.

Although the localization of the CEN2 annuli coincides with the posterior edge of the ISP1 “cap” (Figure 1E and F), depletion of CEN2 from the peripheral annuli did not affect the localization of ISP1 even after 120 h of ATc treatment (Figure S1A, -Shld1/+ATc). Together with a previous finding that the loss of ISP1 has no impact on CEN2 localization (Beck et al., 2010), these data suggest that the assembly and maintenance of the CEN2- and ISP1-containing compartments are independent processes. We then examined the ATc-treated cKD parasites by electron microscopy, to investigate if the depletion of CEN2 from the preconoidal rings and peripheral annuli affected the ultrastructure of the cytoskeletal apical complex. The *CEN2* knockdown parasites (cKD, -Shld1/+ATc) were capable of extruding the conoid (Figure 3D). The structure of the preconoidal rings in the extruded conoid of these parasites also appeared to be normal. However, in slightly under one third of the parasites imaged (4 out of 14), the intra-conoid microtubules were not detectable (Figure 3D, bottom row). This is worth noting because the intra-conoid microtubules were always clearly visible in both wild-type parasites and cKD (+Shld1/-ATc) parasites prepared under the same conditions.

### CEN2 is critical for the parasite lytic cycle

*Toxoplasma* is an obligate intracellular parasite. Its dissemination in the host relies on its ability to progress through multiple rounds of the lytic cycle, which consists of host cell invasion, parasite replication, and egress. To determine the impact of the loss of CEN2 on the *Toxoplasma* lytic cycle, we examined the efficiency of the parasite in forming plaques when *CEN2* is knocked down in cKD parasites using a range of ATc concentrations, with different pre-treatment conditions for two days prior to the plaque assay (+Shld1/-ATc, -Shld1/-ATc, or -Shld1/+ATc) (Figure 4). For all pre-treatment conditions, the cKD parasites did not form plaques when subsequently cultured in the absence of Shld1 with an ATc concentration of 68 nM or higher, while the parental strain (KI:cKD) had comparable plaquing efficiencies under all ATc concentrations tested (0 to 1080 nM). No plaques were observed when the cKD parasites were pre-treated with 270 nM ATc for two days, even when there was no ATc added to the culture medium used in the plaque assay. This suggested CEN2 played an important role in the ability of the parasite to progress through multiple rounds of the lytic cycle.

**Figure 4.**
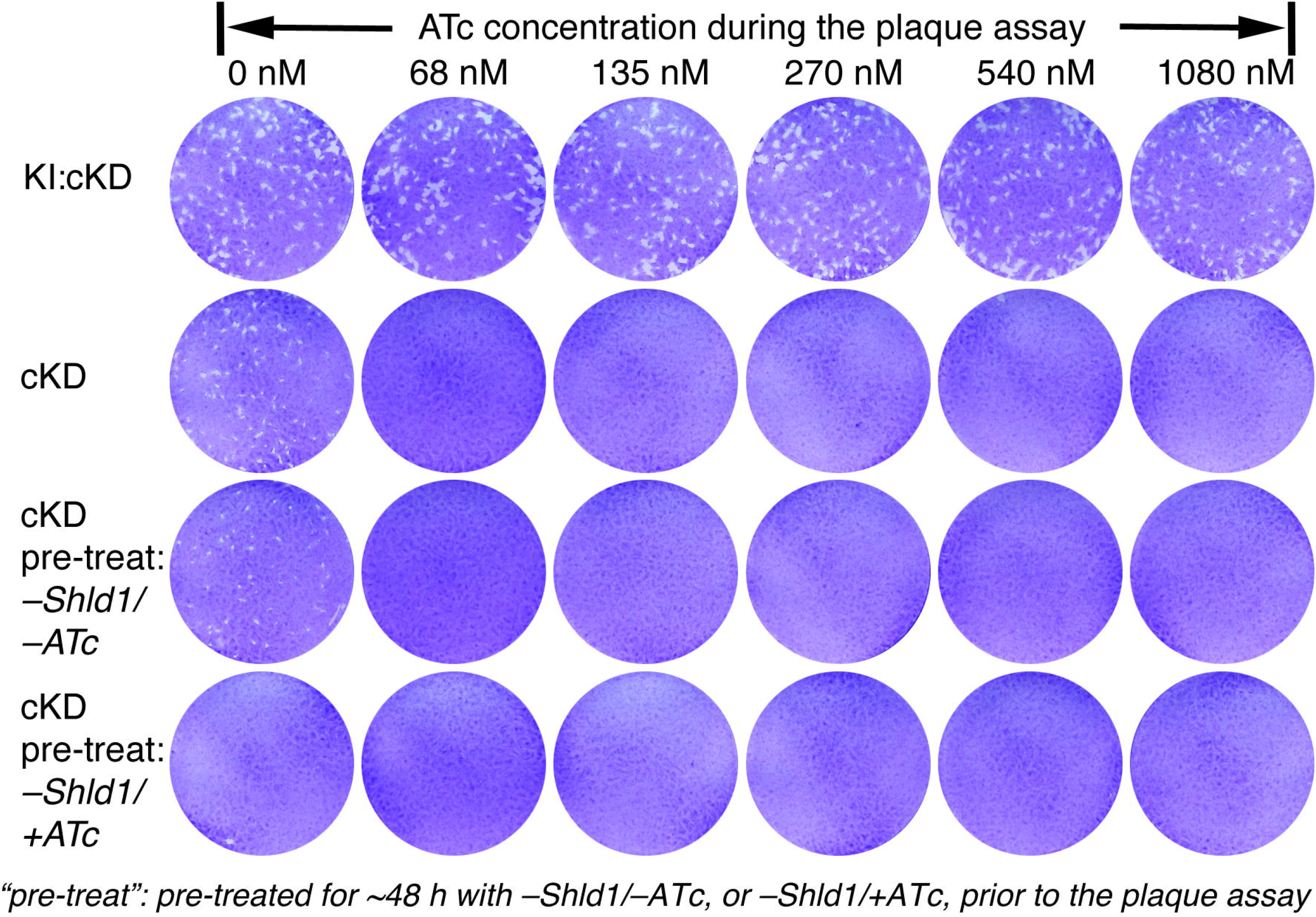
The effect of CEN2 downregulation on the parasite lytic cycle. Plaques formed in the presence of 0, 68, 135, 270, 540 and 1080 nM ATc and without Shld1 by the KI:cKD and cKD parasites. The KI:cKD parasites were maintained in -Shld1/-ATc medium and the cKD parasites were maintained in +Shld1/-ATc medium prior to the plaque assay. “Pre-treat”: The cKD parasites were pre-treated for ~48 h with no drug (-Shld1/-ATc), or 270 nM ATc (-Shld1/+ATc). Each well of HFF monolayers was infected with 100 parasites, grown for 6 days at 37°C, and then fixed and stained with crystal violet. Host cells that remained intact absorbed the crystal violet staining, whereas regions of host cells lysed by the parasites (“plaques”) are clear.

### Parasites deficient in CEN2 are significantly impaired in invasion but not egress

The lytic cycle consists of invasion, replication, and egress. To determine how CEN2 might contribute to the steps of the lytic cycle, we first examined the ability of *CEN2* knockdown parasites to invade host cells, using an established two-color, immunofluorescence-based invasion assay. In this assay, the extracellular (non-invaded) and invaded parasites are differentially labeled due to a difference in antibody accessibility (Carey et al., 2004; Mital and Ward, 2008). Extracellular parasites have the outer surface of their plasma membrane fully exposed, and therefore can be directly labeled by an antibody against a surface antigen (SAG1). On the other hand, parasites that have invaded a host cell can only be labeled after permeabilization of the host cell. We discovered that after ~16 h of -Shld1/+ATc treatment (a timepoint when CEN2 was nearly completely depleted from the preconoidal rings and peripheral annuli, and partially reduced in the centrioles and basal complex), the invasion efficiency of the cKD parasites was ~30% relative to the wild type (P = 0.009, Figure 5A&B and Table 1A). Upon ~48 h of -Shld1/+ATc treatment, invasion by the cKD parasites decreased to ~10% that of the wild-type parasite (P = 0.0014) and ~19% that of the cKD +Shld1/-ATc control (P = 0.005) (Figure 5C&D and Table 1B).

**Figure 5.**
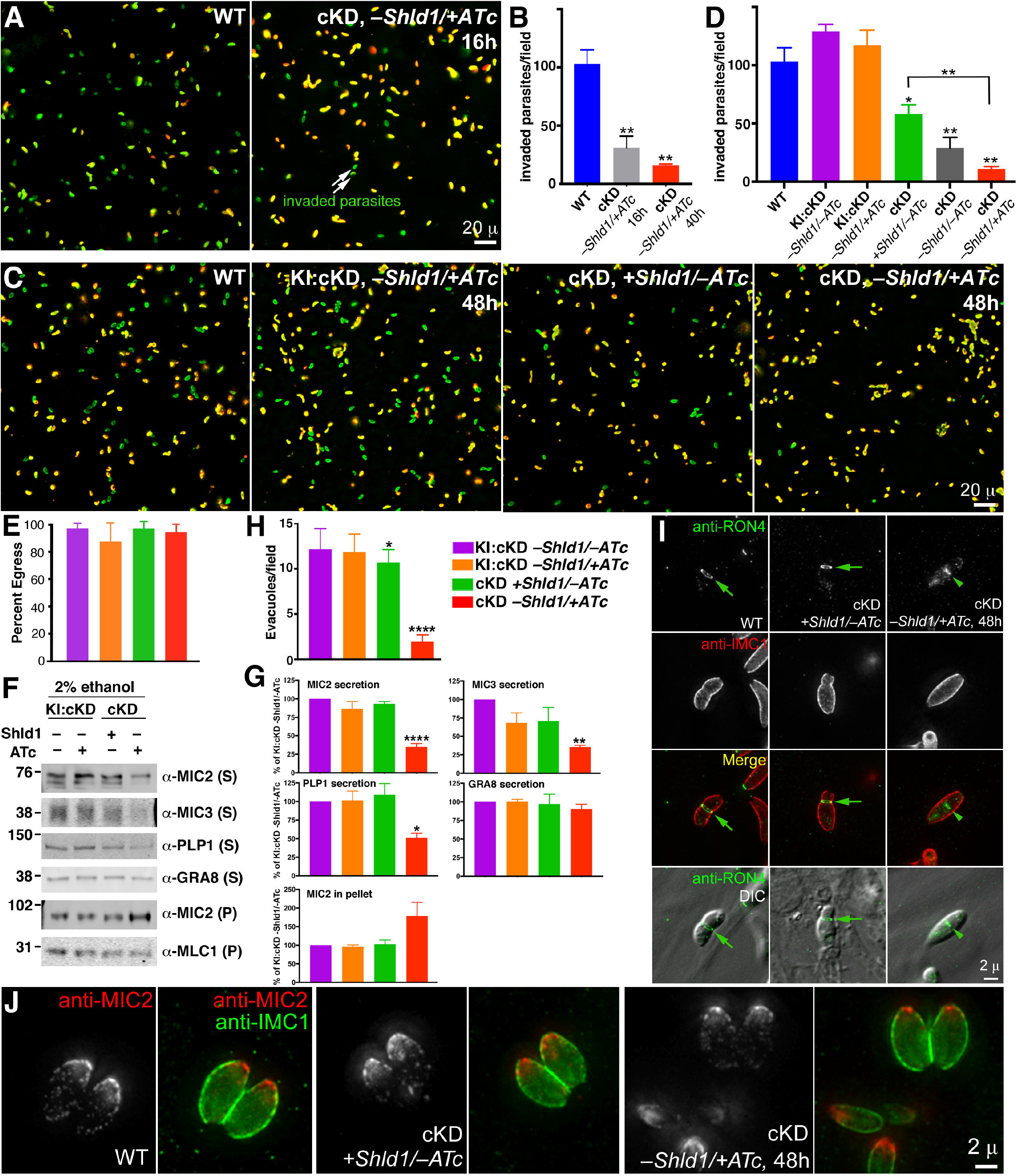
The effect of CEN2 downregulation on parasite invasion, microneme secretion, egress, evacuole formation, and moving junction formation. (A) Representative wide-field epifluorescence images of invasion by RHΔ*hx* parasites (WT), and cKD parasites treated with ATc (-Shld1/+ATc) for 16 h. Parasites that are intracellular (*i.e*., have successfully invaded) are labeled green, and parasites that did not invade are labeled both green and red (*i.e*., appear yellow). Scale bar = 20 μm. (B) Quantification of the mean number of invaded parasites per field for RHΔ*hx* parasites (WT), and cKD parasites treated with ATc (-Shld1/+ATc) for 16 h or 40 h after Shld1 withdrawal (mean + standard error). **: P value <0.01. (C) Representative wide-field epifluorescence images of invasion by RHΔ*hx* parasites (WT), KI:cKD treated with ATc for 48 h, and cKD parasites cultured with Shld1 (+Shld1/-ATc), or treated with ATc (-Shld1/+ATc) for 48 h. Scale bar = 20 μm. (D) Quantification of the mean number of invaded parasites per field for RHΔ*hx* parasites (WT), KI:cKD cultured in -Shld1/-ATc, treated with ATc (-Shld1/+ATc) for 48 h, and cKD parasites cultured with Shld1 (+Shld1/-ATc), no drug (-Shld1/-ATc), or treated with ATc (-Shld1/+ATc) for 48 h (mean + standard error). *: 0.01< P value < 0.05. **: P value 0.01. See also Table 1 for quantification of invasion efficiency relative to WT parasites and the matrix of pairwise P values. (E) Percentage of vacuoles (mean + 2 standard errors) from which parasites activated motility to egress within ~6 min of treatment with 5 μM A23187. For each sample, a total of 12 randomly selected fields in two different dishes were analyzed. The color coding for each condition is as indicated and the same as in (G&H). (F) The unsecreted (pellet, P) and secreted (supernatant, S) fractions of *eGFP-CEN2* knock-in:cKD (KI:cKD, -Shld1/-ATc or -Shld1/+ATc) and cKD (+Shld1/-ATc or -Shld1/+ATc) parasites upon ethanol stimulation, as probed by antibodies against MIC2, MIC3, PLP1, and GRA8, with MLC1 in the pellets as a loading control. The numbers on the left indicate molecular masses in kDa. (G) Levels of MIC2, MIC3, PLP1, and GRA8 in the secreted fractions, and MIC2 in the pellet fractions relative to those from KI:cKD, -Shld1/-ATc. The levels shown for MIC2 in the pellet were normalized to the levels of MLC1 in the pellet for each sample. Error bars: + standard error. *: 0.01< P value <0.05. **: 0.001< P value ≤ 0.01. ****: P value < 0.0001. Results are from at least three independent biological replicates. (H) Mean number of evacuoles observed per field, formed by parasites upon treatment with cytochalasin D. The color coding for each condition is the same as in (E). Error bars: + standard error. *: 0.01< P value <0.05. ****: P value <0.0001. Note that for KI:cKD -Shld1/±ATc, and cKD +Shld1/-ATc, the mean and standard error were calculated from 10 randomly selected fields. For cKD -Shld1/+ATc, the mean and standard error were calculated from 10 fields that had at least one evacuole each. The mean thus overestimates the efficiency of evacuole formation under this condition as most fields did not contain any evacuoles. (I) Wide-field deconvolution images of RHΔ*hx* parasites (WT) and cKD parasites cultured with Shld1 (+Shld1/-ATc), or treated with ATc for 48 h after Shld1 withdrawal (-Shld1/+ATc), in which RON4 (green), and IMC1 (red, a marker for the cortex of mature and daughter parasites) were labeled by immunofluorescence after extracellular parasites were incubated with host cells for a short period of time (*i.e*. “pulse invasion”). Moving junctions (arrows) were readily observed for the WT and the cKD +Shld1/-ATc parasites, but never for CEN2 depleted parasites (cKD, -Shld1/+ATc, 48 h). The RON4 signal (arrowhead) close to the middle of the CEN2 depleted parasite might associate with the Golgi or other organelles in the secretory pathway. Scale bar = 2 μm. (J) Representative wide-field deconvolution images of RHΔ*hx* parasites (WT) and cKD parasites cultured with Shld1 (+Shld1/-ATc), or treated with ATc for 48 h after Shld1 withdrawal (-Shld1/+ATc), in which IMC1 (green) and MIC2 (red) were labeled by immunofluorescence. Scale bar = 2 μm.

**Table 1.**
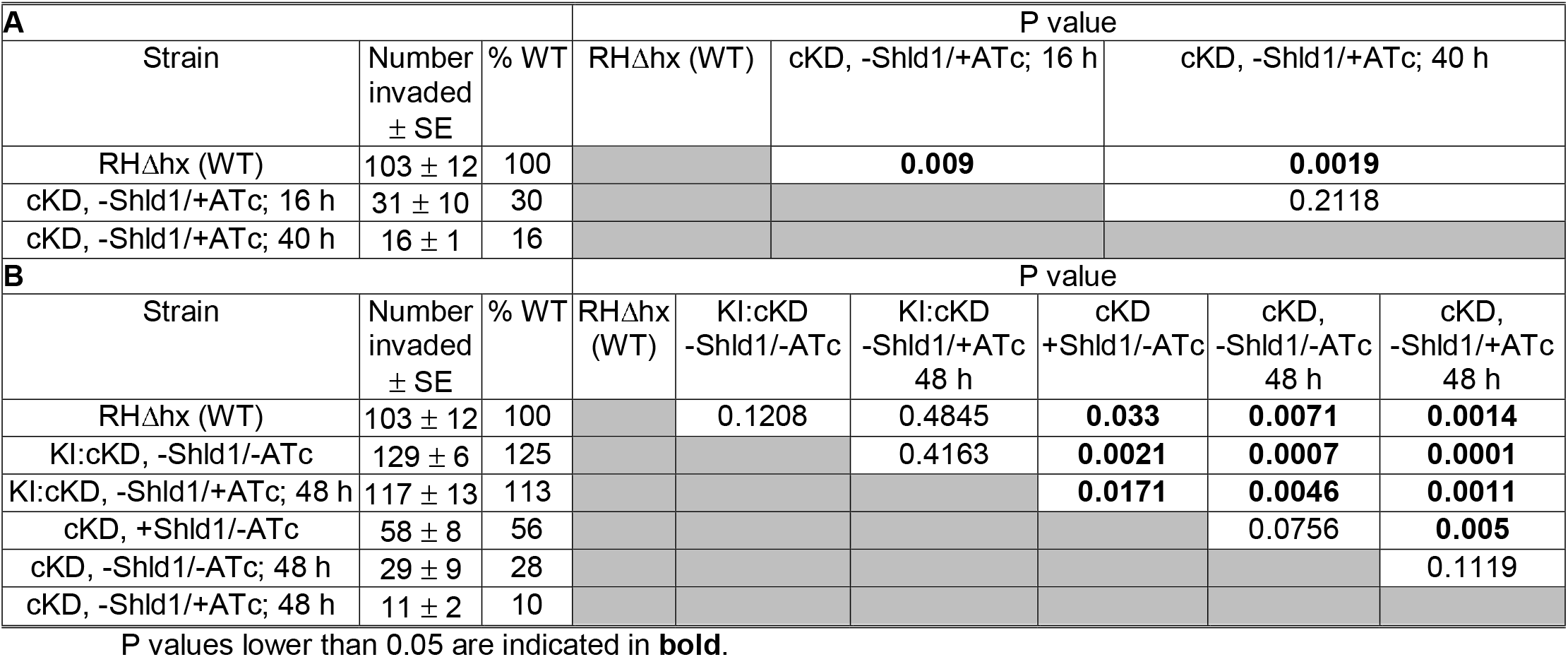
Quantification of invasion [mean number of intracellular parasites per field ± standard error (SE)] by RHΔ*hx* parasites (WT), KI:cKD and cKD parasites with Shld1 (+Shld1/-ATc), no drug (-Shld1/-ATc), or ATc (-Shld1/+ATc). See also Figure 5. The number of intracellular parasites per field was counted in 15 fields per parasite line or treatment, in each of three independent biological replicates. P values from unpaired, two-tailed Student’s *t* tests are indicated on the right.

| A |  |  | P value |  |  |  |  |  |
| --- | --- | --- | --- | --- | --- | --- | --- | --- |
| Strain | Number invaded<br>± SE | % WT | RHΔhx (WT) | cKD, -Shld1/+ATc; 16 h | cKD, -Shld1/+ATc; 40 h |  |  |  |
| RHΔhx (WT) | 103 ± 12 | 100 |  | 0.009 | 0.0019 |  |  |  |
| cKD, -Shld1/+ATc; 16 h | 31 ± 10 | 30 |  |  | 0.2118 |  |  |  |
| cKD, -Shld1/+ATc; 40 h | 16 ± 1 | 16 |  |  |  |  |  |  |
| B |  |  | P value |  |  |  |  |  |
| Strain | Number invaded<br>± SE | % WT | RHΔhx (WT) | KI:cKD<br>-Shld1/-ATc | KI:cKD<br>-Shld1/+ATc<br>48 h | cKD<br>+Shld1/-ATc | cKD,<br>-Shld1/-ATc<br>48 h | cKD,<br>-Shld1/+ATc<br>48 h |
| RHΔhx (WT) | 103 ± 12 | 100 |  | 0.1208 | 0.4845 | 0.033 | 0.0071 | 0.0014 |
| KI:cKD, -Shld1/-ATc | 129 ± 6 | 125 |  |  | 0.4163 | 0.0021 | 0.0007 | 0.0001 |
| KI:cKD, -Shld1/+ATc; 48 h | 117 ± 13 | 113 |  |  |  | 0.0171 | 0.0046 | 0.0011 |
| cKD, +Shld1/-ATc | 58 ± 8 | 56 |  |  |  |  | 0.0756 | 0.005 |
| cKD, -Shld1/-ATc; 48 h | 29 ± 9 | 28 |  |  |  |  |  | 0.1119 |
| cKD, -Shld1/+ATc; 48 h | 11 ± 2 | 10 |  |  |  |  |  |  |

P values lower than 0.05 are indicated in **bold**.

Comparison among the wild-type, parental (KI:cKD), and cKD parasites under various Shld1 and ATc treatment conditions further supports that parasite invasion efficiency is linked to the level of CEN2 expression (Figure 5C&D and Table 1B). Expressing FP-tagged CEN2 from both the endogenous locus as well as the ectopic TetO7Sag4 promoters, the parental line (KI:cKD, -Shld1/-ATc) invades at ~125% of the level of the wild-type parasite (P = 0.1208). Derived from the KI:cKD line by deletion of the *eGFP-CEN2* gene from the endogenous locus, the cKD line invades at a significantly lower efficiency than KI:cKD. Consistent with the differences in the mAppleFP-CEN2 levels (Figure 3B), the cKD parasites invade with different efficiencies under the +Shld1/-ATc (~56% of the wild-type, P = 0.033), -Shld1/-ATc (~28%, P = 0.0071), and -Shld1/+ATc (~10%, P = 0.0014) conditions. The lower level of invasion for the cKD +Shield/-ATc parasites compared to the wild type is most likely because expression of ddFKBP-mAppleFP-CEN2 in the cKD parasites is driven by the TetO7Sag4 promoter, which is known to be weak. Similar differences among the wild-type, parental (KI:cKD) and cKD parasites in TATi based systems have been observed before (Huynh and Carruthers, 2006).

In contrast to the significant defect in invasion, CEN2 depletion did not result in a delay in calcium-induced egress. Upon stimulation with the calcium ionophore A23187, *CEN2* knockdown parasites (cKD, -Shld1/+ATc, ~48 h) in more than 90% of vacuoles activated motility to egress within 6 min, similar to what was observed for the parental line (KI:cKD) (Figure 5E).

### Parasites deficient in CEN2 are significantly impaired in micronemal secretion and form significantly fewer evacuoles and moving junctions

Parasite invasion requires the secretion of adhesins from the micronemes, which allows the parasite to attach to and move into the host cell (Carruthers et al., 1999; Carruthers and Sibley, 1997; Huynh and Carruthers, 2006; Huynh et al., 2003). The adhesins, such as the micronemal protein MIC2, are secreted onto the parasite surface and then cleaved, releasing the ectodomain into the supernatant (Huynh and Carruthers, 2006; Huynh et al., 2003). A deficiency in secretion of these adhesins results in severely impaired host cell invasion. Upon examining the amount of released ectodomain in the supernatant by western blotting, we discovered that after ~40 h of *CEN2* knockdown, ethanol stimulated MIC2 secretion was partially inhibited. This was reflected in the reduced amount of MIC2 in the secreted fraction and a corresponding increase in the pellet fraction (Figure 5F&G). The secretion of two other micronemal proteins, PLP1 and MIC3, was also significantly reduced, indicating that the impact of CEN2 depletion on micronemal secretion is global, and not limited to a specific micronemal protein. In contrast, the secretion of a dense granule protein, GRA8, was not affected by CEN2 depletion (Figure 5F&G).

During parasite invasion, micronemal secretion is followed by protein discharge from the rhoptries, another set of apically located organelles. Rhoptry discharge is typically visualized and assessed by the evacuole assay (Hakansson et al., 2001), in which the parasites are prevented from completing host cell invasion by treatment with cytochalasin D. The *CEN2* knockdown parasites (cKD, -Shld1/+ATc, ~48 h) generated significantly fewer evacuoles compared with the parental line (Figure 5H), consistent with the invasion defect. Secretion from the micronemes and rhoptries also both contribute to the formation of the moving junction, a structure critical for parasite invasion (Alexander et al., 2005; Beck et al., 2014; Lamarque et al., 2011; Mital et al., 2005; Tonkin et al., 2011). This ring-shaped constriction forms during parasite invasion to line the opening of the nascent parasitophorous vacuole (Figure 5I). As the parasite moves forward during invasion, it squeezes through this constriction, and eventually the moving junction caps the basal end of the parasite when invasion is complete. While the moving junction (labeled by anti-RON4) was readily observable for wild-type or cKD +Shld1/-ATc parasites in pulse invasion assays (Figure 5I), we were never able to see a single case of moving junction formation when CEN2 was knocked down (cKD, -Shld1/+ATc, 48 h).

Despite the defect in secretion, CEN2 depletion does not appear to perturb the localization of the secretory organelles in intracellular parasites. Similar to the wild-type parasite, the micronemal vesicles (marked by anti-MIC2 antibody) were concentrated in the apical portion of the parasite when CEN2 was depleted (Figure 5J), indicating that the *CEN2*-containing structures play a role in the mechanics of micronemal secretion rather than the biogenesis or distribution of micronemal organelles. CEN2 depletion also did not affect the distribution of the rhoptries (labeled by anti-ROP2,3,4 and anti-RON2-4 antibodies) and dense granules (labeled by anti-GRA8 antibody) (Figure S1B-G).

### CEN2 knockdown results in abnormal replication patterns

The centrosome (including the centrioles and the spindle pole) is a critical organelle for cell division, and the basal complex has been shown to be involved in parasite cytokinesis (Heaslip et al., 2010). We therefore examined the impact of CEN2 depletion on parasite replication (Figure 6). As the level of CEN2 decreased due to ATc treatment and Shld1 withdrawal, an increasing proportion of vacuoles contained parasites that deviated from the canonical one-to-two division process (Figure 6A). After ~24 h with ATc treatment (Figure 6B & 7A-B), only ~11% of the vacuoles had parasites displaying abnormal division or growth. At this point, CEN2 had been depleted from the preconoidal rings and peripheral annuli to an undetectable level (Figure 3). We thus propose that the CEN2 pools in the preconoidal rings and peripheral annuli do not play a major role in parasite replication. By ~84 h of ATc treatment, ~64% of the vacuoles contained parasites that showed signs of abnormal replication (Figure 6B). In addition to multiple (>2) daughters assembling in mother parasites (Figure 7C), the parasites were often swollen in size and some had undergone nuclear division without cytokinesis, *i.e*., had multiple nuclei but no daughters (Figure 7D). Although the kinetics of CEN2 depletion from the centrioles and the basal complex are similar, these replication defects are most likely due to CEN2 depletion from the centrioles, because the presence and appearance of the basal complex marker IAP1 (Frenal et al., 2014) were not significantly affected (Figure S2A-B). In addition, while centriole duplication occurred in CEN2 depleted parasites, segregation of the centrioles appeared to be perturbed (Figure 7C), as indicated by the unequal distribution of foci that contained high anti-CEN3 signal among the daughter parasites. In some of the daughter parasites, no centriole signal was observed (red arrowheads). In the vacuoles containing parasites with replication defects, there was considerable variation in the size and cell cycle stage of the parasites, indicating that CEN2 depletion does not affect a specific checkpoint and that the cumulative effect of knockdown is stochastic in nature. Interestingly, the partitioning and inheritance of membrane-bound organelles such as the apicoplast (Figure S2C-D), were not severely perturbed. Although the appearance of the apicoplast was more heterogeneous in misshapen *CEN2* knockdown parasites (Figure S2D) when compared to parasites expressing stabilized CEN2 (Figure S2C), almost all of these parasites inherited an apicoplast.

**Figure 6.**
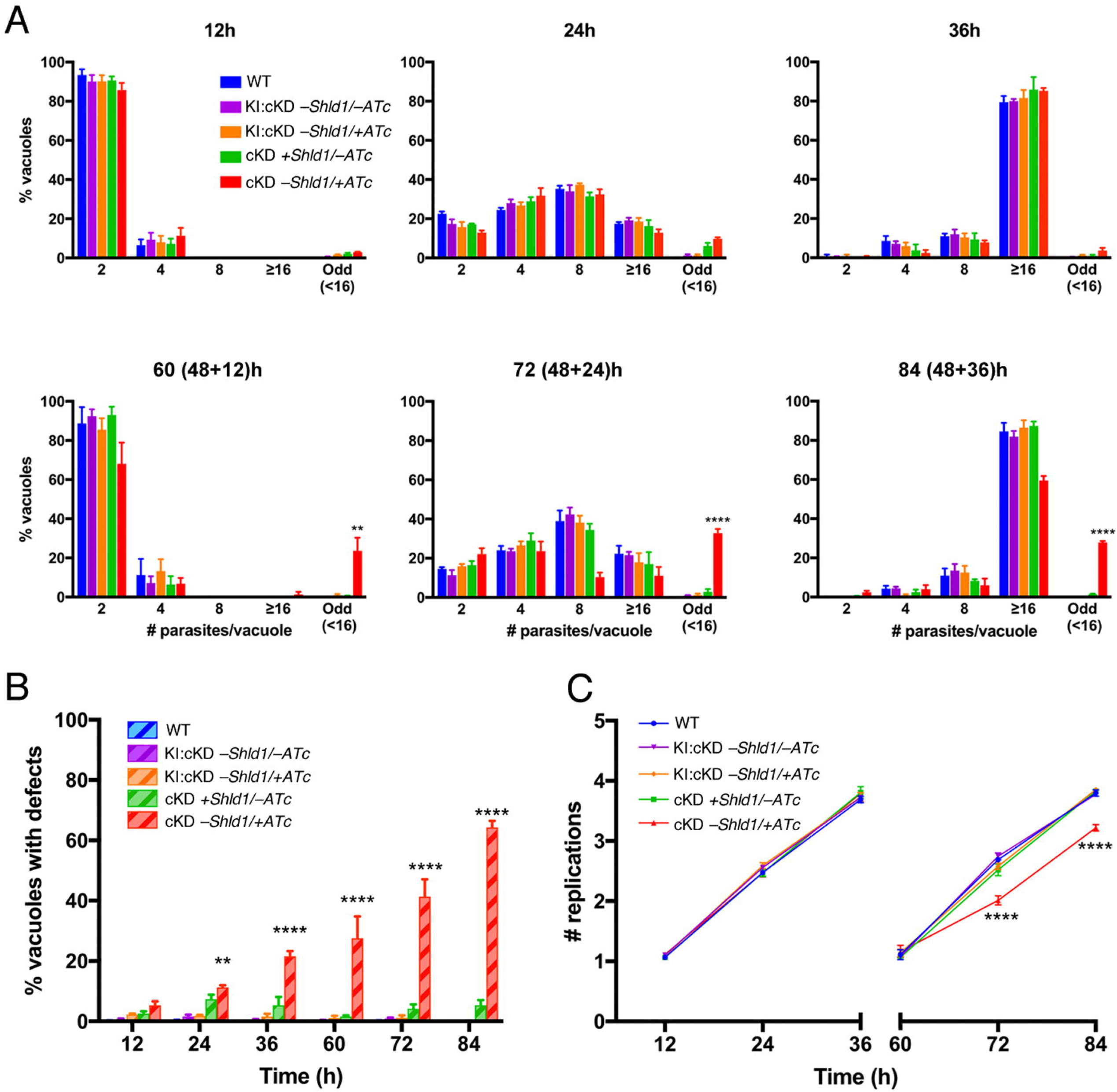
The effect of CEN2 downregulation on parasite replication. (A) Comparison of the intracellular growth of cKD parasites treated with ATc (cKD, -Shld1/+ATc) with that of RHΔ*hx* (WT), KI:cKD (-Shld1/-ATc or -Shld1/+ATc) and cKD parasites cultured with Shld1 (cKD +Shld1/-ATc) parasites at 12, 24, 36, 60 (48+12), 72 (48+24), and 84 (48+36) h after drug removal/addition. To facilitate parasite counting after extended (>48 h) drug treatment times, dishes for the last three timepoints were inoculated with parasites that had been cultured for 48 h in T12.5cm^2^ flasks under the indicated conditions, and allowed to invade a fresh HFF monolayer and proliferate for 12, 24, and 36 h under the corresponding conditions. “Odd”, vacuoles in which the number of parasites was less than 16, and not an integral power of 2. “>16”, vacuoles that contained 16 or more parasites. (B) Percentage of vacuoles containing parasites with replication defects. Phenotypes include an odd number of parasites, enlarged parasites, parasites with multiple nuclei or none at all, and parasites with a single daughter or more than two daughters. (C) Mean replication (doubling) rate. Note that particularly at the 36 h and 84 (48+36) h timepoints, there were many very large vacuoles containing from 16 to upwards of ~128 parasites. However, since the precise number of parasites within each of these large vacuoles could not be calculated, they were grouped into the “16 or more parasites” bin and assigned four doublings (four rounds of replication, equivalent to 16 parasites). Therefore, the replication rates for these timepoints are an underestimate of the actual replication rate. Error bars: standard error. **: 0.001< P value < 0.01. ****: P value < 0.0001 (two-way ANOVA and Bonferroni’s multiple comparisons test), when cKD parasites were compared with RH*Δhx* (WT) or KI:cKD parasites as measured in three independent biological replicates.

**Figure 7.**
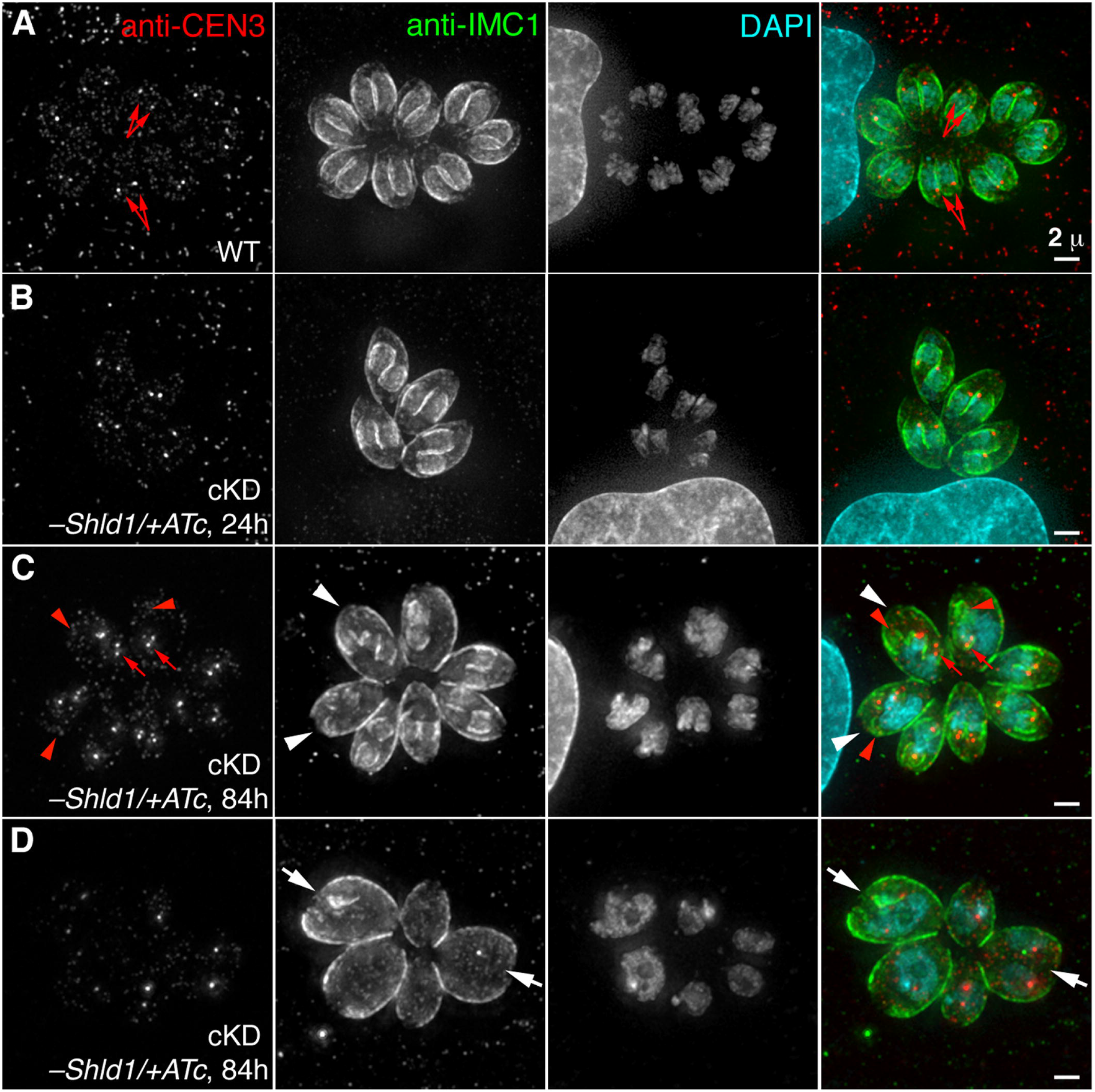
CEN2 knockdown results in abnormal replication patterns. (A) Representative images of untreated RH*Δhx* parasites (WT) at the 24 h timepoint of the replication assays in Figure 6. TgCentrin3 (CEN3), marker for the centrioles (red arrows); TgIMC1 (IMC1), marker for the cortex of mature and daughter parasites; DAPI, fluorescent nucleic acid stain. Scale bar = 2 μm. (B) Images of a vacuole of cKD parasites treated with ATc (-Shld1/+ATc) for 24 h undergoing normal replication. Scale bar = 2 μm. (C&D) Images of one vacuole of cKD parasites treated with ATc (-Shld1/+ATc) for 84 h with more than two daughters forming inside the mother (C, white arrowheads) and another vacuole with parasites that are swollen with large or multiple nuclei (D, white arrows). Notice the uneven distribution of the centrioles (red arrows). Red arrowheads indicate some daughter parasites that receive no centrioles. Scale bars = 2 μm.

## DISCUSSION

In this work, we explore the function of TgCentrin2 (CEN2), a protein that localizes to four different cytoskeletal structures in *Toxoplasma gondii*. Using a dual system that regulates transcription through ATc/TATi and protein stability through Shld1/ddFKBP, we determined the time course of CEN2 depletion by quantitative fluorescence imaging and used it as a guideline to dissect the functional consequences of CEN2 depletion from the individual structures. We found that CEN2 is depleted to an undetectable level from the preconoidal rings and peripheral annuli much earlier than from the centrioles and basal complex. The ordered depletion of CEN2 from the two sets of structures is correlated with the development of defects in two distinct steps of the parasite lytic cycle: invasion and replication. This indicates that CEN2 is recruited to multiple cytoskeletal structures and is required for distinct aspects of parasite physiology.

### The dual-regulation system enables tight control of genes critical for the parasite lytic cycle

Numerous attempts to generate a knockout mutant of *CEN2* using Cre-LoxP and CRISPR-Cas9 based methods failed. Furthermore, a *CEN2* knockdown line in which *CEN2* downregulation solely relied on transcriptional control with ATc quickly became unresponsive to ATc treatment, probably as a result of selecting for mutants that had lost ATc sensitivity within the population. While essentiality cannot be positively proven (*i.e*. essential genes cannot be deleted, but unsuccessful attempts to generate knockout mutants do not prove essentiality), these results suggest that the parasite is highly sensitive to altered *CEN2* expression and that *CEN2* is likely to be essential. By implementing a dual-regulation system that simultaneously utilizes ATc/TATi to control transcription and Shld1/ddFKBP to control protein stability, the expression of *CEN2* can be consistently downregulated. Although growth adaptation can still be detected over long-term passage (>3 months), the phenotype is sufficiently stable for analysis as long as care is taken to use low passage cultures. Given that a single conditional regulation technique (*e.g*. ATc-based transcriptional regulation or ddFKBP-based protein degradation) is always “leaky” to some extent, a knockdown strategy that employs orthogonal methods can be useful for characterizing the functions of genes that are otherwise difficult to manipulate.

### The kinetics of protein knockdown are affected by structural inheritance through cellular replication

In addition to the half-life of transcripts and their protein products, the kinetics of protein knockdown are affected by structural inheritance through cellular replication. Given that *Toxoplasma* and other apicomplexans divide by building daughters inside the mother, if everything else is equal, components of the cortical cytoskeleton or an organelle regenerated *de novo* with every replication should be depleted much faster than those of structures inherited from the mother parasite. For proteins such as CEN2 that are components of multiple structures, this is an important consideration when delineating their functions. For instance, our data indicate that structural inheritance plays an important role in the differential depletion of CEN2 from the four structures: the CEN2 populations in the preconoidal rings and peripheral annuli are depleted much earlier, because these structures are made *de novo* during parasite replication. In contrast, the CEN2 population in the centrioles persists over many generations after downregulation, because each daughter inherits one pair of the duplicated centrioles from the mother. However, factors other than structural inheritance must also contribute to the differential depletion of CEN2, as the CEN2 population in the basal complex does not fit this pattern. A new basal complex is built for each daughter with every generation (Hu, 2008; Hu et al., 2006), but the depletion of CEN2 from the basal complex follows a trend similar to that of the centrioles rather than the preconoidal rings or peripheral annuli. One possible explanation is that as complete CEN2 removal is lethal, only parasites that still express a small amount of CEN2 protein can survive under -Shld1/+ATc conditions. The residual CEN2 protein in these parasites is then preferentially incorporated into the basal complex and centrioles instead of the preconoidal rings and peripheral annuli, perhaps due to differences in binding affinity.

### The role of CEN2 in regulating parasite invasion

The downregulation of CEN2 results in pronounced defects in both parasite invasion and replication. The invasion defect emerged at an early stage of *CEN2* knockdown (~16 h of ATc treatment and Shld1 withdrawal) when CEN2 had disappeared from the preconoidal rings and peripheral annuli and been partially depleted from the centrioles and the basal complex. The timing of depletion suggests that the preconoidal rings and peripheral annuli likely play an important role in parasite invasion. However, the other pools of CEN2 in the centrioles, basal complex, and cytoplasm might also be involved in invasion. In support of this hypothesis, we found that the invasion efficiency of cKD parasites decreased from 30% to 10% upon further CEN2 depletion (~16 h *vs* 48 h of ATc treatment and Shld1 withdrawal) after it had already disappeared from the preconoidal rings and peripheral annuli but was still in the process of being depleted from the centrioles and basal complex. However, it is also possible that further depletion of a very small amount of CEN2 remaining at the apical structures (not detectable by mAppleFP fluorescence) was responsible for the further decrease in invasion efficiency from 16 h to 48 h of CEN2 downregulation.

Part of the invasion defect upon *CEN2* knockdown can be explained by the impaired secretion of MIC2, a major adhesin that mediates parasite attachment to the host cell. As CEN2 is an EF-hand containing protein, it will be of interest to determine whether CEN2 is part of the calcium signaling pathway that controls parasite secretion and invasion. One potential link is *Toxoplasma* phosphoinositide phospholipase C (TgPI-PLC), which is also concentrated in the apical end of the parasite (Fang et al., 2006; Hortua Triana et al., 2018) and has been hypothesized to impact micronemal secretion through regulating calcium homeostasis and the production of phosphatidic acid (Bullen et al., 2016). It is also tempting to speculate that some of the CEN2-containing structures (e.g. the preconoidal rings and peripheral annuli) might be calcium-sensitive contractile structures that gate the release of micronemal contents, but we have not been able to detect specific enrichment of MIC2 at these locations before or after CEN2 depletion. In a subset (~29%) of the *CEN2* knockdown parasites, the intra-conoid microtubules were not detectable. The intra-conoid microtubules have been proposed to serve as tracks for micronemal secretion. It is thus conceivable that structural perturbation of the intra-conoid microtubules interferes with micronemal secretion. However, the lack of intra-conoid microtubule detection in this subset of parasites could also be the result rather than the cause for perturbed micronemal secretion, since the lack of vesicle or protein association might result in decreased stain deposition along the intra-conoid microtubules in electron microscopy.

It is important to note that secretion of the three micronemal proteins tested (MIC2, MIC3, and PLP1) is not completely blocked by the depletion of CEN2, and the invasion defect of the *CEN2* knockdown is more severe than that of the knockout mutants for individual micronemal proteins (Cerede et al., 2005; Gras et al., 2017; Kafsack et al., 2009). Furthermore, the cKD parasites cultured in +Shld1/-ATc had lower invasion efficiency compared to the parental line (KI:cKD), but secreted MIC2 at a similar level. Therefore, CEN2 downregulation must affect other processes that also contribute to invasion. One such process might be moving junction formation, which requires protein secretion from both the micronemes and the rhoptries (Alexander et al., 2005; Beck et al., 2014; Lamarque et al., 2011; Mital et al., 2005; Tonkin et al., 2011). Indeed, we found that *CEN2* knockdown parasites generated significantly fewer evacuoles and moving junctions compared with the wild-type and cKD +Shld1/-ATc parasites. Furthermore, similar to the *CEN2* knockdown parasite, mutants of moving junction components also have low invasion efficiency, but no obvious egress defect (Beck et al., 2014; Mital et al., 2005). It is therefore possible that CEN2 controls parasite invasion partially by regulating moving junction formation. While this is an appealing hypothesis, we should point out that rhoptry secretion occurs after the parasites make contact and form an intimate attachment with the host cell. Therefore, the apparent deficiency in rhoptry discharge and moving junction formation might instead be a consequence of upstream invasion processes disrupted by *CEN2* knockdown rather than the direct cause of the invasion defect.

### The role of CEN2 in regulating parasite replication

CEN2 is a component of the centrioles and the depletion of CEN2 from the centrioles correlates with an increase in abnormal parasite replication. The centrosome (including the centrioles and the spindle pole) has been hypothesized to orchestrate daughter construction and organelle biogenesis in *Toxoplasma* (Dhara et al., 2017; Francia et al., 2012; Hu et al., 2002; Nishi et al., 2008; Suvorova et al., 2015). For instance, knockdown of the centrosome-associated SFAs impairs the initiation of construction of the daughter cytoskeleton (Francia et al., 2012). When a temperature sensitive mutant of MAPKL was inactivated at the non-permissive temperature, the parasite generated an abnormally high number of daughters without completion of cytokinesis (Suvorova et al., 2015). Significant depletion of CEN2 from the centrioles results in a heterogeneous population of multi-nucleated (without daughters), multi-daughter (>2), as well as normal looking parasites. This suggests that CEN2 does not control a specific “checkpoint” or coupling point of the regulatory circuit of the cell cycle. However, it is worth noting that in the parasites that survive prolonged (~120 h) ATc treatment, the CEN2 signal in the centrioles remains detectable, likely because complete CEN2 removal is lethal. Therefore, the consequence of total loss of CEN2 for parasite replication remains unknown. It would also be of interest to determine the functions of other centrins homologs (CEN1 and 3) in the centrioles, and whether they have roles in replication that overlap with CEN2.

The CEN2 containing structures are critical for parasite invasion, replication and survival. Future identification and characterization of structure-specific proteins will facilitate more precise function designation, and aid in the discovery of promising “druggable” targets that lead to multiple points of vulnerability in the parasite.

## MATERIALS AND METHODS

### T. gondii, host cell cultures, and parasite transfection

Tachyzoite *T. gondii* parasites were maintained by serial passage in confluent human foreskin fibroblast (HFFs, ATCC# SCRC-1041) monolayers in Dulbecco’s Modified Eagle’s Medium (DMEM, Life Technologies-Gibco, Cat# 10569-010), supplemented with 1% (v/v) heat-inactivated cosmic calf serum (Hyclone, Cat# SH30087) as previously described (Leung et al., 2017; Liu et al., 2016; Roos et al., 1994). African green monkey renal epithelial cells (BS-C-1, ATCC# CCL-26) used for the invasion assays and rat aortic smooth muscle cells (A7r5, ATCC# CRL-1444) used for live imaging and immunofluorescence assays were cultured in the same manner as HFFs. The *ddFKBP-mAppleFP-CEN2* cKD (“cKD”) parasites were cultured in the presence of 125 nM Shield-1 (a kind gift from Dr. Tom Wandless, Stanford University, Stanford) (Banaszynski et al., 2006) to stabilize its ddFKBP-mAppleFP-CEN2. Downregulation of ddFKBP-mAppleFP-CEN2 was achieved by removal of Shld1 and incubation in medium supplemented with 270 nM anhydrotetracycline (ATc) (Clontech) unless noted otherwise. *T. gondii* transfections were carried out as previously described (Liu et al., 2013).

### Alignment analysis of selected centrin homologs

Protein sequences for selected centrin homologs were aligned using the MUSCLE program accessed through JalView (v2.8.1, http://www.jalview.org) with default parameters and displayed using the Clustal X color scheme.

### Cloning of plasmids

Primers used in cloning DNA fragments and for sequencing are listed in Table S1. Genomic DNA (gDNA) fragments were amplified using gDNA template prepared from RH, RHΔ*hx* or RHΔ*ku80*Δ*hx* (“RHΔ*ku80*” parasites (a kind gift from Dr. Vern Carruthers, University of Michigan, Ann Arbor, MI) (Fox et al., 2009; Huynh and Carruthers, 2009) using the Wizard Genomic DNA Purification Kit (Cat# A1120, Promega, Madison, WI) according to the manufacturer’s instructions. Similarly, coding sequences (CDS) were amplified using *T. gondii* complementary DNA (cDNA). All DNA fragments generated by PCR were confirmed by sequencing.

#### pTKO2_II-eGFP-CEN2 (for generation of *eGFP-CEN2* knock-in parasites)

This plasmid was constructed in the pTKO2_II plasmid backbone (Heaslip et al., 2010) designed for replacement of genes in *T. gondii* by homologous recombination. Three fragments were generated and ligated with the pTKO2_II vector backbone using the corresponding sites. The 3’UTR of *CEN2* (TgGT1_250340) was amplified using S1 and AS1 as primers and RH gDNA as the PCR template, and ligated via the *Nhe*I and *Apa*I sites. The 5’UTR was amplified using S2 and AS2 as primers and RH gDNA as the PCR template, and ligated via the *Not*I and *EcoR*I sites. The *eGFP-CEN2* CDS with a Kozak sequence and flanking *Pme*I and *Rsr*II sites was synthesized (GenScript Inc, NJ) and ligated via the *Pme*I and *Rsr*II sites.

#### pmin-mCherryFP-CEN2 (intermediate plasmid for cloning pTATi1-TetO7Sag4-ddFKBP-mAppleFP-CEN2, below)

This plasmid was constructed by Biomeans Inc (Sugar Land, TX). Briefly, the CEN2 CDS was amplified using primers S3 and AS3 and cloned into pCR-Blunt-II-TOPO (Invitrogen), prior to ligation with the plasmid backbone of pmin-mCherryFP-TgICMAP1 via the *Bam*HI and *Afl*II sites. [pmin-mCherryFP-TgICMAP1 has the same vector backbone as pmin-eGFP-TgICMAP1 (Heaslip et al., 2009) but with mCherryFP in place of eGFP.]

#### pTATi1-TetO7Sag4-ddFKBP-mAppleFP-CEN2 (for generation of *eGFP-TgCEN2* knock-in:cKD and cKD parasites)

To generate pTATi1-TetO7Sag4-ddFKBP-mAppleFP-CEN2, two synthesized DNA or plasmid digestion fragments were generated and ligated with the pTATi1-TetO7Sag4 vector backbone using the corresponding sites. The ddFKBP-mAppleFP fragment features a *Mfe*I restriction site in between the ddFKBP and mAppleFP coding sequences, and was synthesized (GenScript Inc, NJ) and ligated via flanking *Nhe*I-*Bam*HI sites to the *Nhe*I-*Bgl*II sites on the vector backbone. The *CEN2* fragment was derived from the pmin-mCherryFP-CEN2 plasmid and ligated via the flanking *Bam*HI-*Afl*II sites to the *Bgl*II-*Afl*II sites on the vector backbone. The pTATi1-TetO7Sag4 vector backbone was constructed from a four-component HiFi assembly (Cat# E5520, New England Biolabs): 1) the vector backbone, which contains the bacterial origin of replication, beta-lactamase gene for ampicillin resistance, Cre recombinase CDS, and DHFR 3’UTR, was generated from an *Apa*I and *Bmt*I double restriction digest of the plasmid ptub1200bp-mAppleFP-Cre (see below); 2) the TATi1 CDS expression cassette was amplified by PCR using primers S4 and AS4 with the plasmid ptub8-TATi1-HX (a kind gift from Dr. Dominique Soldati-Favre, University of Geneva, Geneva, Switzerland) (Meissner et al., 2002) as the template; 3) the sagCATsag chloramphenicol transferase expression cassette was released from the plasmid ptub1200bp-mAppleFP-Cre by digestion with *Ppu*MI and *Hin*dIII; 4) the ATc-responsive TetO7Sag4 mini-promoter was amplified by PCR using primers S5 and AS5 with the plasmid TetO7Sag4_mycGFP (a kind gift from Dr. Dominique Soldati-Favre, University of Geneva, Geneva, Switzerland) (Meissner et al., 2002) as the template. The ptub1200bp-mAppleFP-Cre plasmid intermediate was constructed by ligating via the *PpuM*I and *Nhe*I sites a truncated tubulin promoter amplified by PCR using primers S6 and AS6 with the plasmid ptub-mAppleFP-TLAP2 (Liu et al., 2016) as the template, and ligating via the *Bgl*II and *Afl*II sites the plasmid backbone ptub-mAppleFP-TLAP2 with the Cre recombinase CDS, which was amplified by PCR using primers S7 and AS7 with the plasmid ptub-Cre-EGFP (Heaslip et al., 2010) as the template.

#### pTKO2_II_mCherryFP-Cre-EGFP (for excision of the *LoxP*-flanked *eGFP-TgCEN2* knock-in expression cassette)

This plasmid was constructed by ligating the Cre-EGFP expression cassette from pmin-Cre-EGFP (Heaslip et al., 2010) with the plasmid backbone of pTKO2_II_mCherryFP (Liu et al., 2013) via the *Not*I and *Apa*I sites.

#### *Generation of knock-in, conditional knockdown, and transgenic parasites eGFP-CEN2* knock-in parasites

Approximately 1 x 10^7^ RH*Δhx* parasites were electroporated with 50 μg of pTKO2_II-eGFP-CEN2 linearized with *Not*I and selected with 25 μg/mL mycophenolic acid and 50 μg/mL xanthine for four passages, and enriched by FACS for parasites with an intermediate level of eGFP signal to reduce the number of parasites in the population that had not undergone double homologous recombination. Clones were screened by fluorescence light microscopy, and confirmed by diagnostic gDNA PCRs to have the *CEN2* endogenous locus replaced by a *LoxP*-flanked eGFP-CEN2 expression cassette. One clone was further verified by Southern blotting; this clone was used for the subsequent generation of *eGFP-TgCEN2* knock-in:cKD and cKD parasites.

#### *eGFP-TgCEN2* knock-in:cKD (KI:cKD) parasites

The *eGFP-CEN2* knock-in parasites were transfected with 30 μg of pTATi1-TetO7Sag4-ddFKBP-mAppleFP-CEN2 that subsequently integrated multiple times, randomly into the parasite genome after selection with 20 μM chloramphenicol.

#### *ddFKBP-mAppleFP-CEN2* cKD (cKD) parasites

KI:cKD parasites (~1 x 10^7^) were electroporated with 20 μg of pTKO2-mCherryFP-Cre-eGFP to excise the knock-in expression cassette between the two *LoxP* sites, selected with 80 μg/mL of 6-thioxanthine for two passages, and screened for the loss of eGFP-CEN2 fluorescence. Clones were confirmed by diagnostic gDNA PCRs. One clone (clone 10) was further verified by Southern blotting and used in all of the experiments reported here. In the resultant parasite line (cKD), the expression level of ddFKBP-mAppleFP-CEN2 is controlled by ATc and Shld1. The parasite line was maintained in the presence of 125 nM Shld1 unless indicated otherwise.

### Generation of the rat TgCEN3 and TgIAP1 antibodies

Purified recombinant TgCEN3 (TogoA.00877.a.A1.PS00788) and TgIAP1 (TogoA.17172.a.A1.PW29285) proteins (kind gifts from the Seattle Structural Genomics Center for Infectious Disease, Seattle, WA) were used to inject rats for antibody production (Cocalico Biologicals, Inc) and sera of the immunized animals were harvested for performing the immunofluorescence labeling of TgCEN3 and TgIAP1.

### FACS and parasite cloning

Fluorescence activated cell sorting was performed using an AriaII flow cytometer (BD Biosciences, San Jose, CA) driven by FACSDiva software at the Indiana University Bloomington Flow Cytometry Core Facility (Bloomington, IN). To subclone parasites by limiting dilution, 3-25 parasites (depending on the survival rate of specific parasite lines after sorting) with the desired fluorescence profile were sorted per well of a 96-well plate containing confluent HFF monolayers. Wells were screened 7-9 days after sorting for single plaques.

### Southern blotting

Southern blotting was performed as previously described (Liu et al., 2016; Liu et al., 2013) with probes synthesized using components based on the NEBlot Phototope Kit (New England BioLabs, Cat# N7550) and detected using components based on the Phototope-Star detection kit (New England BioLabs, Cat# N7020). All *T. gondii* gDNA was prepared from freshly egressed parasites and extracted using the Wizard Genomic DNA Purification kit (Cat# A1120, Promega, Madison, WI).

To probe and detect changes at the *CEN2* genomic locus in the RHΔ*hx* (WT), *eGFP-CEN2* knock-in (KI), KI:cKD, and cKD parasites, 5 μg of gDNA was digested with *Sca*I. A CDS probe (282 bp) specific for Exon 1 of *CEN2* was amplified from *T. gondii* RHΔ*hx* gDNA using primers S8 and AS8, and used as a template in probe synthesis. A probe specific for the region upstream of the *CEN2* genomic locus (168 bp) was amplified from plasmid pTKO2_II-eGFP-CEN2 using primers S9 and AS9 and used as a template in probe synthesis.

### Plaque assay

Plaque assays were performed as previously described with some modifications (Liu et al., 2016). A total of 100, 200 or 500 freshly egressed, pre-treated parasites (see below) was added to each well of a 12-well plate containing a confluent HFF monolayer, and incubated undisturbed for 6-7 days. The pre-treatment medium and corresponding incubation medium were either the regular parasite growth medium, or the medium supplemented with the concentrations of either ATc or Shld1 as indicated in the text and figure legends. Infected monolayers were then fixed with 3.7% (v/v) formaldehyde at 25°C for 10 min, washed with Dulbecco’s phosphate-buffered saline (DPBS), stained with 2% (w/v) crystal violet, 20% (v/v) methanol in DPBS for 15 min, gently rinsed with distilled water and air dried.

### Wide-field deconvolution microscopy

Image stacks were acquired at 37°C using a DeltaVision imaging station (GE Healthcare / Applied Precision) fitted onto an Olympus IX-70 inverted microscope base. A 60X silicone oil immersion lens (Olympus 60X U-Apo N, NA 1.3) used with or without an auxiliary magnification of 1.5X, or 100X oil immersion lens (Olympus 100X UPLS Apo, NA = 1.40) with immersion oil at a refractive index of 1.524 was used for imaging. 3D image stacks were collected with a z-spacing of 0.3 μm unless otherwise noted. Images were deconvolved using the point spread functions and software supplied by the manufacturer. All samples for wide-field deconvolution and for 3D-SIM (below) were prepared in 35-mm dishes (#1.5) with a 20- or 14-mm microwell (P35G-1.5-20-C or P35G-1.5-14-C; MatTek) in phenol red-free, CO_2_-independent medium for live imaging and in DPBS with 10 mM sodium azide for fixed samples. Contrast levels were adjusted to optimize the display.

### Three-dimensional structured-illumination microscopy (3D-SIM)

3D-SIM image stacks were acquired using an OMX imaging station (GE Healthcare / Applied Precision, Seattle, WA). A 100X oil immersion lens (NA = 1.40) with immersion oil at a refractive index of 1.516 or 1.518 was used; stacks were collected with a z-spacing of 0.125 μm. Images were deconvolved using the point spread functions and software supplied by the manufacturer.

### Electron microscopy

Negative staining of whole mount, detergent-extracted parasites was performed as described in (Leung et al., 2017). To determine the effect of CEN2 depletion on the parasite cytoskeleton, cKD parasites were treated with -Shld1/+ATc for 144 h prior to processing for EM analysis.

### *Quantification of mAppleFP-CEN2 fluorescence in* cKD parasites

#### a. Parasite culture and imaging conditions

*T. gondii* cultures were grown in culture medium +Shld1/-ATc (125 nM Shld1), -Shld1/-ATc, or -Shld1/+ATc (270 nM ATc) for 12, 24, 48, 72, 96 or 120 h prior to live imaging. The average intensity of individual CEN2-containing structures in cKD parasites cultured in +Shld1/-ATc was used to calculate the baseline fluorescence. Multiple flasks of parasites were coordinated with staggered timing such that a freshly lysed flask was used to inoculate 35-mm dishes (#1.5) with a 20-mm microwell (P35G-1.5-20-C; MatTek) approximately 16 h prior to imaging. This yielded smaller vacuoles containing 2, 4 or 8 parasites to facilitate imaging each of the four CEN2-containing compartments within the parasite for the quantification of mAppleFP-TgCEN2 signal. The medium in each dish was replaced with pre-warmed, phenol red-free CO_2_-independent medium (custom order, SKU#RR060058; Gibco/Life Technologies) prior to imaging. Dishes were imaged in a humidified environmental chamber maintained at 37°C.

Twenty full fields of view (1024 x 1024 pixels) were collected with 2×2 binning for quantitative analysis. For each field of view, a reference differential interference contrast (DIC) image was acquired followed by a 3D stack of three z-slices, spaced 1.0 μm apart, of fluorescence images in the mAppleFP and eGFP channels. Irradiation of each field of view was minimized to prevent photobleaching and kept consistent to reduce variability in the quantification.

#### b. Image analysis

Image analysis strategies and procedures used here were carried out as previously described in (Murray, 2017). The Semper software package [source code kindly provided by Dr. Owen Saxton (Murray-Edwards College, University of Cambridge, United Kingdom) (Saxton et al., 1979)] was used for image analysis.

#### c. Grouping of integrated fluorescence and statistical analysis

A minimum of twenty vacuoles were quantified per parasite line and drug treatment. Vacuoles containing a smaller number of intracellular parasites (2, 4 or 8 parasites) were used for quantification, and the mAppleFP-CEN2 signal was classified into four groups corresponding to each of the compartments it localizes to in the parasite (preconoidal rings, peripheral annuli, centrioles, and basal complex). All photons/s measurements shown are per individual structure, per parasite except for the peripheral annuli, which were quantified collectively (*i.e*., all annuli in one parasite were grouped as a single value). Since the basal complexes of parasites tend to cluster within a vacuole, these were quantified collectively for each vacuole and subsequently calculated per parasite, based on the number of parasites observed in the corresponding reference DIC image. Data were analyzed using two-way ANOVA and Dunnett’s multiple comparisons tests with GraphPad Prism v7 (La Jolla, CA).

### Immunofluorescence assay for intracellular parasites

*T. gondii*-infected HFF monolayers growing in a 3.5 cm glass-bottom dish were fixed with 3.7% (v/v) formaldehyde in DPBS for 10 min, permeabilized with 0.5% (v/v) Triton X-100 (TX-100) in DPBS for 15 min, and blocked in 1% (w/v) BSA in DPBS for 30-60 min, followed by antibody labeling (see below). Dishes were incubated with primary antibodies for 30-60 min followed by incubation with secondary antibodies for 30-60 min unless otherwise noted. Primary antibodies and dilutions used were as follows: rabbit anti-ACP, 1:500 (a kind gift from Dr. David Roos, University of Pennsylvania, Philadelphia, PA and Dr. Geoff McFadden, University of Melbourne, Melbourne, Australia) (Waller et al., 1998); rabbit anti-TgIMC1, 1:1,000 (a kind gift from Dr. Con Beckers, University of North Carolina, Chapel Hill) (Mann and Beckers, 2001); mouse anti-TgISP1, 1:1,000 (a kind gift from Dr. Peter Bradley, University of California, Los Angeles) (Beck et al., 2010); mouse anti-TgMIC2 6D10, 1:1,000 (a kind gift from Dr. Vern Carruthers, University of Michigan, Ann Arbor) (Carruthers et al., 2000); mouse monoclonal anti-GRA8 and mouse monoclonal anti-IMC1 45.36 antibodies, 1:1,000 (kind gifts from Dr. Gary Ward, University of Vermont, Burlington) (Carey et al., 2000; Ward and Carey, 1999); mouse anti-ROP2,3,4, 1:1,000 (a kind gift from Dr. Jean-François Dubremetz, Université de Montpellier, Montpellier, France) (Leriche and Dubremetz, 1991); rabbit anti-RON2-4 and rabbit anti-RON4, 1:500 (kind gifts from Dr. Maryse Lebrun, Université de Montpellier, Montpellier, France) (Lamarque et al., 2011; Lebrun et al., 2005); rat anti-CEN3, 1:1,000 (this study); and rat anti-IAP1, 1:1,000 (this study). Secondary antibodies, all used at 1:1,000 dilution were: goat anti-rabbit IgG Alexa488, 1:1,000 (Cat#A11034, Molecular Probes); goat anti-rabbit IgG Alexa568 (Cat#A11036, Molecular Probes); goat anti-rat IgG Alexa488 (Cat#A11006, Molecular Probes); goat anti-rat IgG Alexa568 (Cat#A11077, Molecular Probes); goat anti-mouse IgG Cy3 (Cat#115-166-003, Jackson ImmunoResearch); goat anti-mouse IgG Alexa488 (Cat# A11029, Molecular Probes); and goat anti-mouse IgG Alexa568 (Cat# A11031, Molecular Probes). To label the nucleus, 4’,6-diamidino-2-phenylindole (DAPI) was used at a final concentration of 1 μg/mL. All immunofluorescence labeling steps were performed at room temperature.

### Invasion assays

Invasion assays were performed as previously described (Leung et al., 2017) with the following modifications. Since we observed during routine culturing that there was a clear invasion defect for a subset of the parasite lines, preparation of the cultures was adjusted in terms of the amount of inoculum, and timed such that all parasite preparations would be at the same stage at the time of harvest. For ATc treatment (-Shld1/+ATc), at approximately 48, 40 or 16 h prior to harvest, the medium for the indicated subset of parasites was changed to that containing 270 nM ATc and no Shld1.

For the first round of immunolabeling, the primary antibody was mouse anti-SAG1 (Argene, Cat# 11-132; 1:1,000 dilution for 30 min), followed by goat anti-mouse Alexa568 (1:1,000 dilution for 30 min). For the second round of immunolabeling, the primary antibody was rabbit anti-SAG1 (a kind gift from Dr. Lloyd Kasper, Dartmouth College, Lebanon, NH; 1:1,000 dilution for 30 min), followed by goat anti-rabbit IgG Alexa488 (1:1,000 dilution for 30 min). The dishes were imaged at low magnification (Olympus 20X UPlanApo, numerical aperture (NA) = 0.70) for a total of 15 full-fields of view per sample for each of three independent experiments. Fields were randomly selected using the Alexa488 channel. The mean number of parasites counted per sample, per replicate was ~3,926.

Semi-automated quantification of invaded parasites was performed using FIJI [ImageJ v. 2.0.0-rc-65/1.51s; (Schindelin et al., 2012; Schneider et al., 2012)] as previously described (Leung et al., 2017). Pairwise comparisons were made with unpaired, two-tailed Student’s t-tests using GraphPad Prism, v7 (GraphPad, La Jolla, CA).

### Microneme secretion (excretory/secretory antigen assay)

Microneme secretion assays were performed as previously described (Leung et al., 2017) with some modifications. Freshly egressed parasites were harvested and resuspended in DMEM supplemented with 1% (v/v) cosmic calf serum (Hyclone, Cat# SH30087.3). Excreted/secreted antigen preparation was performed by incubating 1 x 10^8^ parasites (adjusting the final volume to 500 μL) at 37°C for 7-15 min in 2% (v/v) ethanol to assess induced secretion. Tubes were placed on ice for 5 min immediately afterwards, and centrifuged at 1,000 x *g* for 6 min at 4°C. 450 μL of the supernatant was removed and centrifuged again, and 400 μL of the second supernatant was concentrated ~tenfold using an Amicon Ultra 3,000 MWCO centrifugal filter device (Millipore) before adding an equal volume of 4X NuPAGE sample buffer. The pellet was washed with DPBS and resuspended in 50 μL RIPA buffer (150 mM NaCl, 1% (v/v) NP-40, 0.5% (v/v) sodium deoxycholate, 0.1% (w/v) SDS, 50 mM Tris, pH 7.4), and then treated with benzonase nuclease (Santa Cruz Biotechnology, TX) for 15 min at 37°C before the addition of an equal volume of 4X NuPAGE sample buffer. Samples were incubated at 75°C for 10 min and resolved using NuPAGE 4-12% Bis-Tris gels, and blotted by wet transfer to nitrocellulose membrane. Membranes were processed for western blotting by probing with mouse mAb anti-TgMIC2 6D10, 1:5,000; mouse mAb anti-TgMIC3 T4 2F3, 1:200 (a kind gift from Dr. Maryse Lebrun, Université de Montpellier, Montpellier, France) (Cerede et al., 2005); rabbit anti-TgPLP1, 1:1,000 (a kind gift from Dr. Vern Carruthers, University of Michigan, Ann Arbor) (Kafsack et al., 2009); mouse anti-GRA8 (1:5,000), and rabbit anti-TgMLC1 (1:2,000; a kind gift from Dr. Con Beckers, University of North Carolina, Chapel Hill) (Gaskins et al., 2004) in TBS-T with 5% (w/v) BSA or blocking solution (1% (w/v) casein, Hammarsten, 0.1 M maleate pH 7.5, 0.1 M NaCl), washed with TBS-T, followed by goat anti-mouse IRDye 680RD, goat anti-mouse IRDye 800CW, or goat anti-rabbit IRDye 800CW infrared dye-conjugated antibodies (1:10,000; LI-COR) in TBS-T with 5% (w/v) nonfat dry milk or blocking solution. Blots were scanned in the 700- and 800-nm channels using an LI-COR Odyssey Classic imaging system (LI-COR Biosciences, Lincoln, NE), and band intensities were plotted and then quantified with local background subtraction using the Gel Analyzer tool in FIJI (ImageJ v. 2.0.0-rc-65/1.51s).

### Egress assay

Calcium induced egress assays were performed as previously described (Heaslip et al., 2011). KI:cKD parasites were maintained in -Shld1/-ATc medium and cKD parasites were cultured in +Shld1/-ATc medium. For ATc treatment (-Shld1/+ATc), KI:cKD and cKD parasites were treated with 270 nM ATc and no Shld1 for ~46-52 h, prior to the egress assay.

### Evacuole assay

Evacuole assays were performed as previously described (Hakansson et al., 2001) except the parasites were harvested and resuspended in ENDO buffer (44.7 mM K_2_SO_4_, 10 mM MgSO_4_, 106 mM sucrose, 5 mM glucose, 20 mM Tris-H_2_SO_4_, 3.5 mg/mL BSA, adjusted to pH 8.2) (Endo and Yagita, 1990) prior to induction of evacuole formation with 1 μM cytochalasin D in DMEM medium supplemented with 1% (v/v) cosmic calf serum. KI:cKD parasites were cultured in -Shld1/-ATc medium and cKD parasites were cultured in +Shld1/-ATc medium. For ATc treatment (-Shld1/+ATc), KI:cKD and cKD parasites were treated with 270 nM ATc and no Shld1 for ~48 h prior to the evacuole assay. Anti-ROP2,3,4 antibody was used for visualizing rhoptry discharge. Note that for KI:cKD -Shld1/±ATc, and cKD +Shld1/-ATc, the mean and standard errors were calculated from 10 randomly selected fields. For cKD -Shld1/+ATc, the mean and standard error were calculated from 10 fields that had at least one evacuole each. The mean thus overestimates the efficiency of evacuole formation under this condition as most fields did not contain any evacuoles.

### Pulse invasion assay for analyzing moving junction formation

Pulse invasion assays were performed as previously described (Parussini et al., 2012) with some modifications. For ATc treatment (-Shld1/+ATc), cKD parasites were treated with 270 nM ATc and no Shld1 for 48 h prior to the pulse invasion assay. Parasites were harvested from large vacuoles and resuspended in ENDO (invasion non-permissive) buffer pre-warmed to 37°C, added to a ~90% confluent HFF monolayer in a 3.5 cm glass-bottom dish, and incubated at 37°C for 30 min. The ENDO buffer was gently exchanged with invasion permissive medium (DMEM + 1% (v/v) heat-inactivated cosmic calf serum) pre-warmed to 37°C. RHΔ*hx* (wild-type) parasites and cKD parasites maintained in +Shld1/-ATc medium were allowed to invade at 37°C for 1 min 30 s; cKD parasites treated with ATc (-Shld1/+ATc) were allowed to invade for 1 min 30 s, 2 min 30 s, 4 min, 8 min, and 18 min. The dishes were washed once with DPBS prior to immunofluorescence labelling as described above.

### Intracellular replication assay

Intracellular replication assays were performed as previously described (Heaslip et al., 2010), except parasites were grown for 12, 24, 36, 60, 72 or 84 h in medium with no drug supplementation (-Shld1/-ATc), with 125 nM Shld1 (+Shld1/-ATc) or with 270 nM ATc (-Shld1/+ATc). To ensure that the number of parasites in a vacuole was countable, for the last three timepoints (*i.e*., 60, 72 and 84 h), parasites were first cultured for 48 h in T12.5cm^2^ flasks in the indicated conditions (-Shld1/-ATc, +Shld1/-ATc, or -Shld1/+ATc), released by mechanical disruption of the host cell monolayer, then used to inoculate dishes containing the same medium conditions as the originating flasks, for an additional 12, 24 or 36 h. At each timepoint, dishes were processed for immunofluorescence as described above, using antibodies for markers of the mother and daughter cortex (TgIMC1), centrioles (TgCEN3), basal complex (TgIAP1) and a fluorescent nucleic acid stain (DAPI) to count the number of parasites per vacuole and to assess the percentage of replication defects. The number of parasites per vacuole was determined as 2, 4, 8, ≥16 or “odd” (*i.e*., where the number of parasites was less than 16, and not an integral power of 2). For Figure 6B, vacuoles with any of the following phenotypes were classified as having replication defects: an odd number of parasites, enlarged parasites, parasites with multiple nuclei or none at all, and parasites with a single daughter or more than two daughters. Note that for Figure 6C, particularly at the 36 h and 84 h timepoints, there were many very large vacuoles containing from 16 to upwards of ~128 parasites. However, since the precise number of parasites within each of these vacuoles could not be calculated, they were grouped into the “16 or more parasites” bin and assigned four doublings (four rounds of replication, equivalent to 16 parasites). Therefore, the replication rates for these timepoints are an underestimate of the actual replication rate.

For every timepoint, a minimum of 200 vacuoles was assessed for each parasite line and condition in each of three independent biological replicates, except for the cKD parasites treated with ATc, in which 100 vacuoles were quantified per timepoint because its invasion deficiency resulted in many fewer vacuoles formed. Data were analyzed using two-way ANOVA and Bonferroni’s multiple comparisons tests with GraphPad Prism v7 (La Jolla, CA). Where statistically significant, multiplicity adjusted P values for comparisons are indicated with asterisks.

## ACKNOWLEDGEMENTS

We thank Dr. Owen Saxton (Murray-Edwards College, University of Cambridge, United Kingdom) for the Semper image processing software, Drs. Con Beckers (University of North Carolina, Chapel Hill) for the rabbit anti-TgIMC1 and anti-TgMLC1 antibodies, Peter Bradley (University of California, Los Angeles) for the mouse anti-TgISP1 antibody, Vern Carruthers (University of Michigan, Ann Arbor) for the mouse mAb 6D10 anti-TgMIC2 antibody and rabbit anti-TgPLP1 antibodies, Jean-François Dubremetz (Université de Montpellier, Montpellier, France) for the mouse anti-ROP2,3,4 antibody, Lloyd Kasper (Dartmouth College, Lebanon, NH) for the rabbit anti-TgSAG1 antibody, Maryse Lebrun (Université de Montpellier, Montpellier, France) for the rabbit anti-RON2-4, rabbit anti-RON4, and mouse anti-TgMIC3 antibodies, David Roos (University of Pennsylvania, Philadelphia, PA) and Geoff McFadden (University of Melbourne, Melbourne, Australia) for the rabbit anti-ACP antibody, Gary Ward (University of Vermont, Burlington) for the mouse anti-GRA8 and mouse anti-IMC1 antibodies, Richard Day (Indiana University School of Medicine, Indianapolis, IN) for the pmAppleFP-C1 plasmid, Dominique Soldati-Favre (University of Geneva, Geneva, Switzerland) for the TetO7Sag4_mycGFP and ptub8-TATi1-HX plasmids, Tom Wandless (Stanford University, Stanford, CA) for Shield-1, and the Seattle Structural Genomics Center for Infectious Disease (Seattle, WA) for the purified recombinant TgCEN3 and TgIAP1. We thank Christiane Hassel of the Indiana University Bloomington (IUB) Flow Cytometry Core Facility for assistance with flow cytometry, and Dr. James Powers of the IUB Light Microscopy Imaging Facility for assistance and support with light microscopy (NIH S10-RR028697). We would also like to thank Dr. Amanda Rollins and Qing Zhang for technical support. This study was supported by American Heart Association Postdoctoral Fellowships (16POST31330004 and 18POST34090005) awarded to J.M.L., and funding from the March of Dimes (6-FY18-674), the National Institutes of Health/National Institute of Allergy and Infectious Diseases (R01-AI132463) awarded to K.H., and facility funding from the Indiana Clinical and Translational Sciences Institute to K.H., funded in part by Grant UL1 TR001108 from a National Institutes of Health, National Center for Advancing Translational Sciences, Clinical and Translational Sciences Award.

