## Supplementary_Figures_Table for "TgCentrin2 is required for invasion and replication in the human parasite *Toxoplasma gondii*"

**Figure S1.** CEN2 depletion does not affect the distribution of ISP1, rhoptries, or dense granules. (A) Images of cKD parasites treated for 120 h with ATc (-Shld1/+ATc), labeled with antibodies for ISP1 (red), a marker for the apical cap, and IMC1 (green), a marker for the cortex of mature and daughter parasites. Scale bar = 2 µm.

(B-D) Images of RHΔ*hx* parasites (WT, B), and cKD parasites cultured with Shld1 (+Shld1/-ATc, C), or treated for 48 h with ATc (-Shld1/+ATc, D), labeled with antibodies for the rhoptries. Red: anti-RON2-4, a marker for the rhoptry neck. Green: anti-ROP2,3,4, markers for the rhoptry bulb. Scale bars = 2 µm.

(E-G) Images of RHΔ*hx* parasites (WT, E), and cKD parasites cultured with Shld1 (+Shld1/-ATc, F), or treated for 48 h with ATc (-Shld1/+ATc, G), labeled with antibodies for GRA8 (red), a marker for the dense granules, and IMC1 (green). Scale bars = 2 µm.

**Figure S2.** CEN2 depletion does not have a major impact on construction of the basal complex or inheritance of the apicoplast.
Representative images of cKD parasites cultured with Shld1 (+Shld1/-ATc, A&C) or treated with ATc (-Shld1/+ATc) for 84 h (B) or 87 h (D). The parasites were labeled with antibodies for IMC1 (green), and IAP1 (A&B, red), a marker for the basal complex, or acyl carrier protein (ACP, C&D, red), a marker for the apicoplast. Scale bars = 2 µm.

**Table S1.** Primers used in this study.

### Figure S1

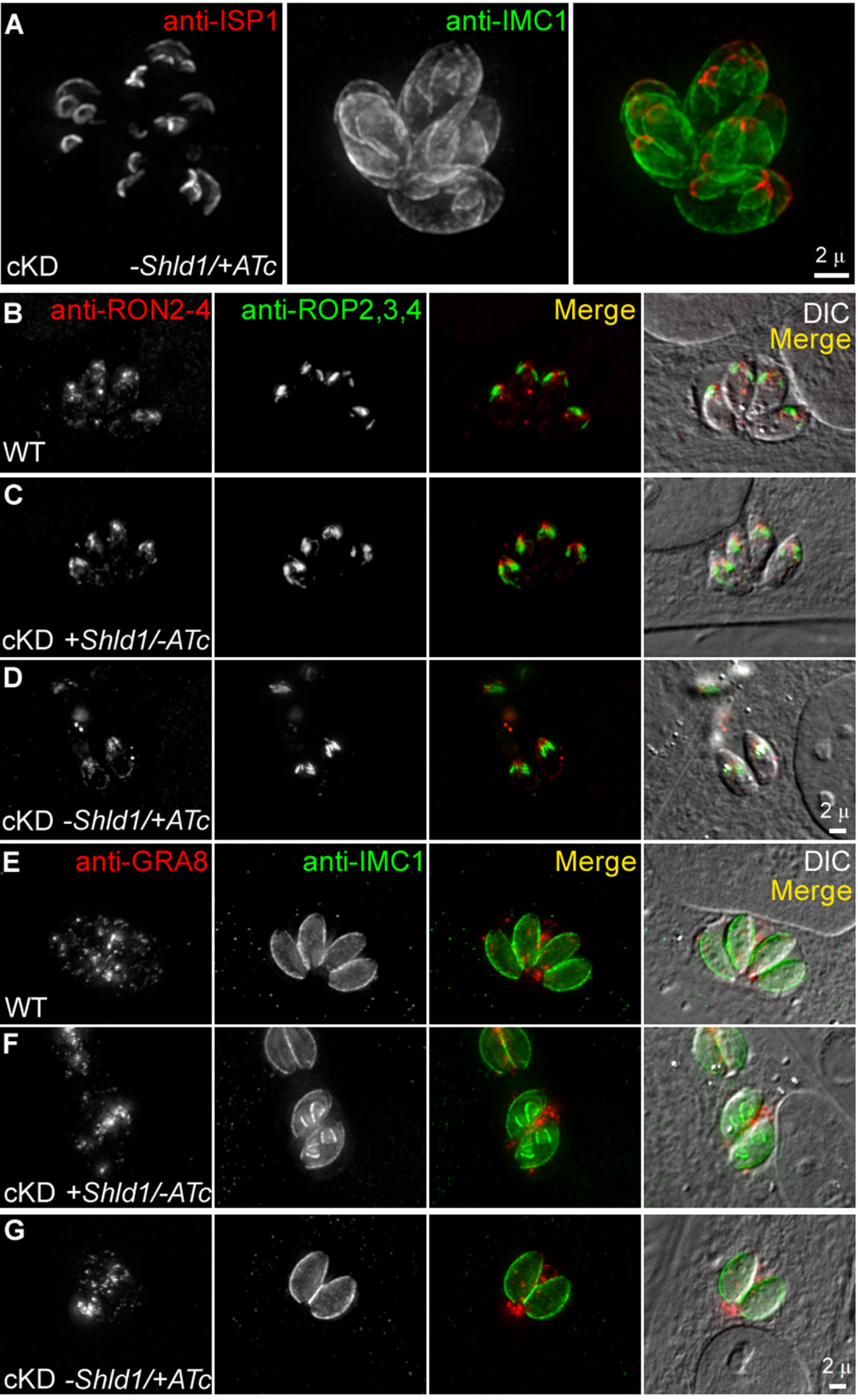
A
anti-ISP1
anti-IMC1
cKD
-Shld1/+ATc
2 μ
B
anti-RON2-4
anti-ROP2,3,4
Merge
DIC
Merge
WT
C
cKD +Shld1/-ATc
D
cKD -Shld1/+ATc
2 μ
E
anti-GRA8
anti-IMC1
Merge
DIC
Merge
WT
F
cKD +Shld1/-ATc
G
cKD -Shld1/+ATc
2 μ

### Figure S2

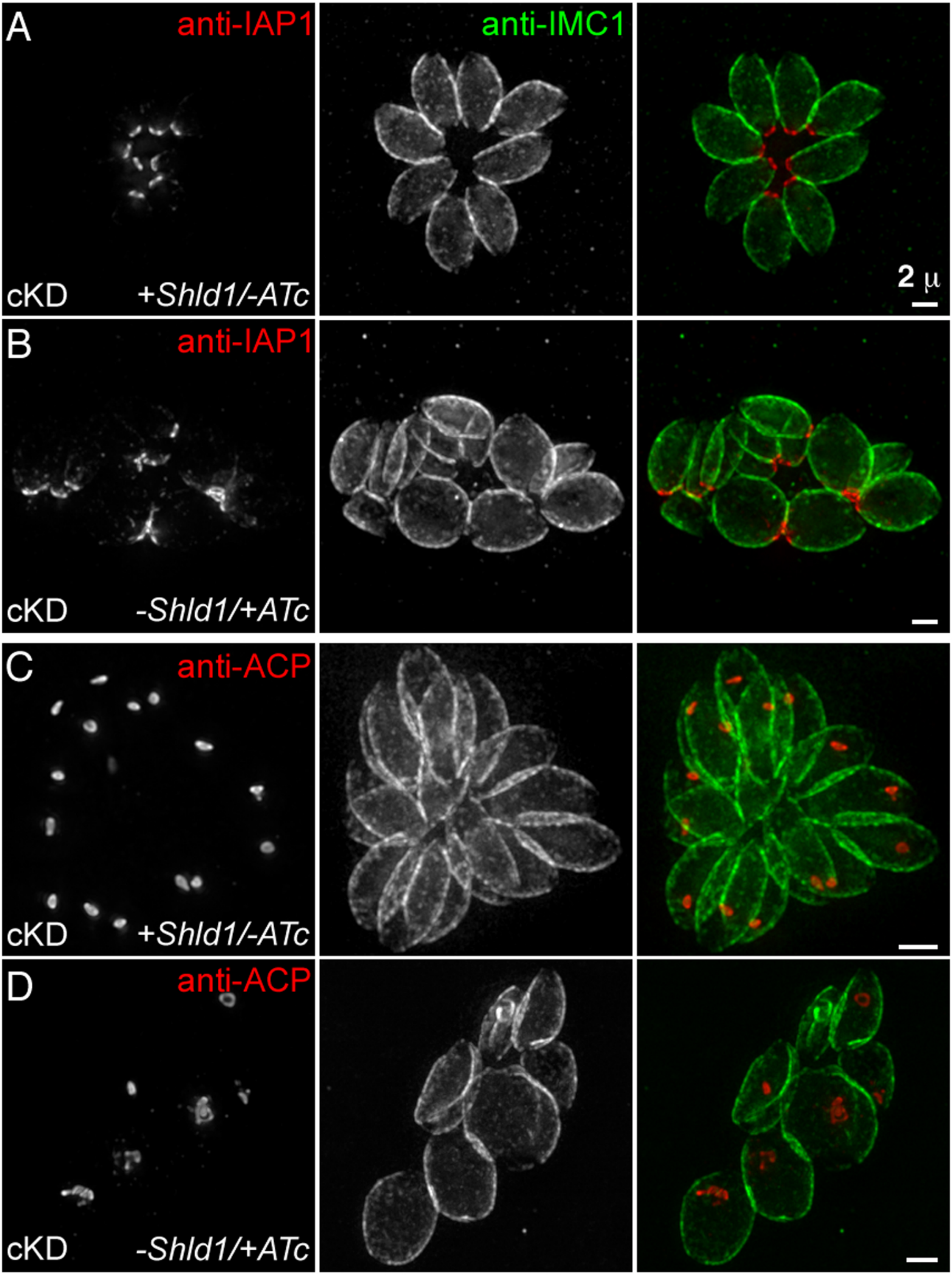

A
anti-IAP1
anti-IMC1
cKD
+*Shld1/-ATc*
2 μ
B
anti-IAP1
cKD
*-Shld1/+ATc*
C
anti-ACP
cKD
+*Shld1/-ATc*
D
anti-ACP
cKD
*-Shld1/+ATc*

**Table S1. Primers used in this study.**

| Name | Sequence (5' to 3') |
|---|---|
| S1 | ACTGGCTAGCCAAGGCTGTCGATTCAACAGAGAGC |
| S2 | AGTCGCGGCCGCCTCGCACTTTTCGCAGGGCATCTTG |
| S3 | CGATGGATCCCAGCGAGGAGCACTGCGAGGGGCGAG |
| S4 | CTAAAGGGAACAAAAGCTGGGTACCGGTACCGGGCCCCCCCTCG |
| S5 | CTCCGGCTTGCAACCAAGGACCCGTAATACGACTCACTATAGGGC |
| S6 | ATCGAGGACCCCTGATGAACTTGGCTTATTCAT |
| S7 | GCGCAGATCTGCCAATTTACTGACCGTACACC |
| S8 | CAGCGAGGAGCACTGCGAGG |
| S9 | GATCGCTTCTCGGTTCCTACCCTG |
| AS1 | ACTGGGGCCCCCTGTGCCCCAAAATGTACCGGAGGC |
| AS2 | ACTGGAATTCGCTCGACAAAAAAAGGCCAAATGTA |
| AS3 | ACGTCTTAAGTCACGGGAAAGTCTTCTTGGTCATGATCG |
| AS4 | CTGCAGGAATTCGATATCAAGCTTAACCGGTTCGACTAAAACAAC |
| AS5 | CGCCCTTGCTCACCATTTTGCTAGCTTTGTCGAAAAAGGGAATTCG |
| AS6 | ATCGGCTAGCGGATCTAAAAGGGAAT |
| AS7 | GCGCCTTAAGCTAGGTGGCGACCGGTCCATCGCCAT |
| AS8 | CAGCTTCGCGGTAATGGCGT |
| AS9 | CCTGTTACGAACGCAAAGATGTGT |
